# Individualized fMRI neuromodulation enhances visuospatial perception: a guided approach targeted towards the neuro-rehabilitation of cortical blindness and deceleration of subjective cognitive impairment

**DOI:** 10.1101/2024.03.03.582390

**Authors:** Anthony Allam, Vincent Allam, Sandy Reddy, Ankit Patel, Emmanouil Froudarakis, Eric M. Rohren, Sameer A. Sheth, T. Dorina Papageorgiou

## Abstract

Neuromodulation is a growing precision-medicine approach to modulating neural activity that can be used to treat neuropsychiatric, and general pathophysiologic conditions. We developed individualized fMRI neuromodulation (iNM) to study the mechanisms of visuospatial perception modulation with the long-term goal of applying it in low-vision patient populations having cortical blindness or visuospatial impairment preceding subjective cognitive impairment. To determine these mechanisms, we developed a direction and coherence discrimination task to engage visual perception (VP), visual imagery (VI), selective extero-intero-ceptive attention (SEIA), and motor planning (MP) networks. Participants discriminated up and down direction, at full and subthreshold coherence under iNM or control (no iNM). We determined the blood-oxygen-level-dependent (BOLD) magnitude as area under the curve (AUC) for VI, SEIA, and MP encoded networks and used a decoder to predict the stimulus from brain maps. The increased AUC BOLD magnitude under iNM across directions and coherences ranged from: 48-76% for SEIA, 26-59% for MP, 20-47% for VI, and 100% for strong VP coherences, but decreased for weak coherences. iNM increased classification performance. Our results imply a causal role of iNM-induced visuospatial mechanisms in strengthening these networks and provide a pathway for more accurate encoding models and effective treatment.

## Introduction

Neuromodulation is a rapidly expanding precision-medicine approach that includes a spectrum of invasive and non-invasive interventions ranging from invasive deep brain stimulation to non-invasive transcranial magnetic stimulation, transcutaneous electrical nerve stimulation, and fMRI neurofeedback. Each of these interventions modulate neural activity, and can be used to treat psychiatric, neurological, and general pathophysiological conditions such as cerebral visual impairment (CVI; Pamir et al., 2021), specifically the visuospatial perception.(Cho et al., 2015; D. Papageorgiou et al., 2014). CVI refers to damage in retrochiasmatic pathways and surrounding cortical visual areas that presents with visual deficits such as perception of complex motion, characterized by difficulty navigating the environment (McDowell & Dutton, 2019).

In this study, we developed and applied a guided, *i*ndividualized fMRI closed-loop *<u>n</u>*euro*<u>m</u>*odulation (iNM; Figure 1A) approach by targeting (with 1 mm precision) and strengthening each participant’s unique anatomy and functional extents and intensity of the middle temporal (MT/V5, also known as the hMT+ complex) that includes the medial superior temporal (MST) area. The rationale to target these areas is based on macrostimulation studies in macaques, which have shown that MT and MST signals guide visual motion perception (Braddick, 1974; Ditterich et al., 2003; Liu & Newsome, 2005a, 2005b; Nichols & Newsome, 2002; Price & Born, 2013; Tootell et al., 1995; Zaksas & Pasternak, 2006a).

**Figure 1.**
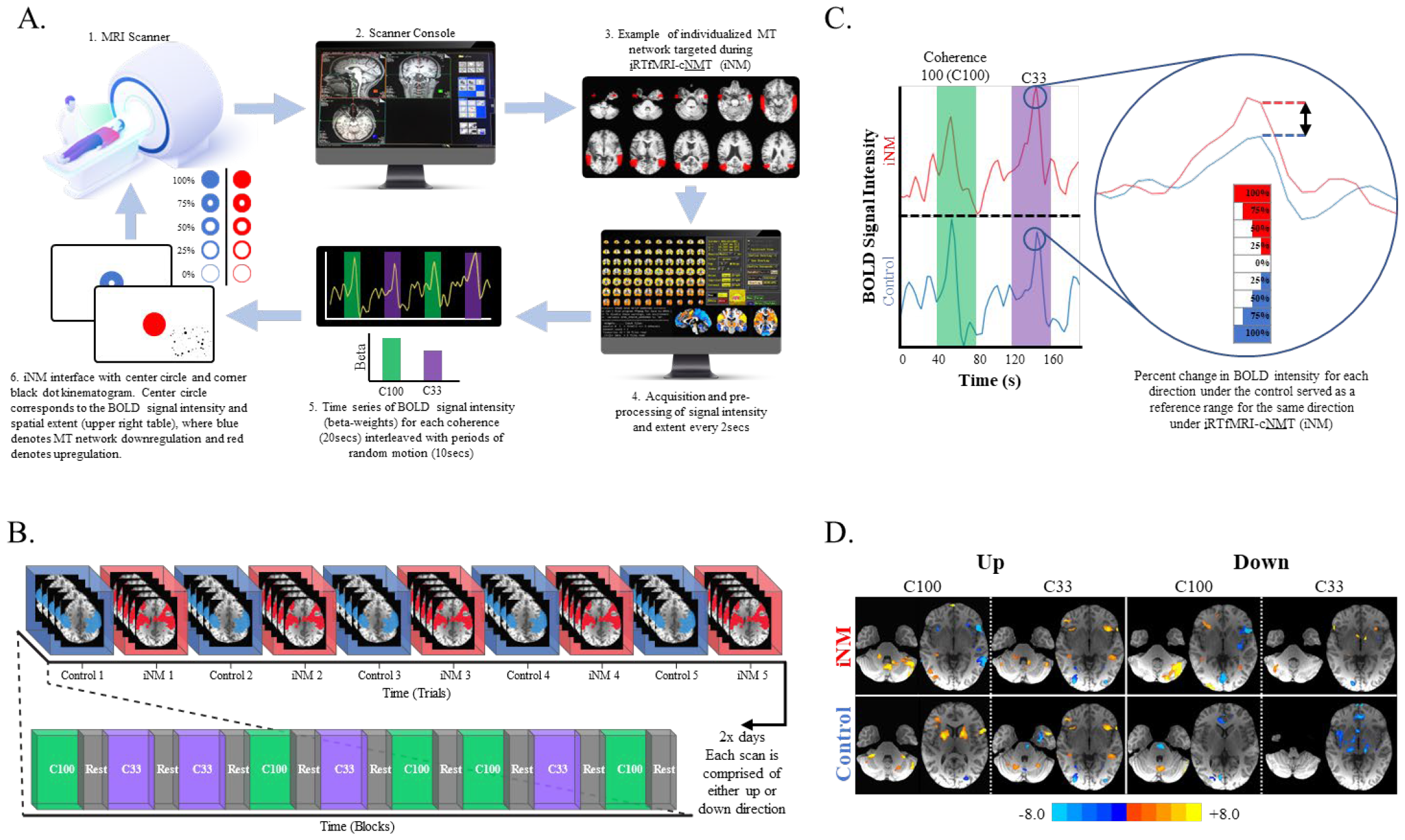
Individualized real-time functional MRI closed-loop neuromodulation (iNM) Intervention. **A. iNM strengthens visual perception and visual imagery networks**: 1) and 2) High-resolution anatomical images were acquired and registered to the Siemens’ console computer; 3) Each participant’s individualized MT and MST networks were delineated and contoured; 4) iNM data extracted from individualized networks (**Figure SI-9**) was preprocessed and general linear modeling was used to decode each coherence level in real-time every TR=2 sec (yellow line denotes the time series during green, purple, and black periods representing up or down direction at coherence level C100, C33, and baseline-random motion period, respectively); 5) The BOLD signal intensity and spatial extent (individualized network) for each direction was computed via a GLM, and beta weights were updated, see Fig. 1B. 6) iNM interface shows the extent of the circle filled, which directly corresponds to the percent upregulation (red fill) or downregulation (blue fill) of the BOLD signal intensity and spatial extent. **B. Task design (details under Figure SI-2a and 2b):** The study was performed over two days in which 5 control and 5 iNM runs were completed in an alternating fashion. Each run was performed in a temporal sequence that included: a coherent motion block lasting 20 seconds interleaved with a 10 second baseline-random motion block. The level of coherence was counterbalanced across runs, and blocks ensuring that the same coherence levels were not stacked back-to-back. **C. Computation of the iNM signal:** The neuromodulation stimulus was calculated as a percent change of the BOLD signal intensity during the task compared to the baseline-random motion block during each control run which served as a reference. If the percent BOLD signal change during a 2-sec interval was higher than the 10th percentile, the iNM interface indicated a 10% upregulation, and if less than the 10th percentile, the iNM interface indicated downregulation. **D. Brain activation maps show increased activation in the iNM condition compared to control:** GLM-generated activation maps for each coherence level and motion direction in iNM (top) and control-no cNMT (bottom) conditions are shown; voxelwise p-value = 0.05, cluster size > 20 voxels.

The long-term goal of our iNM approach is two-fold: 1) to neurorehabilitate visual perception (VP) following cortical blindness; and 2) to decelerate visuospatial impairment, as a prodromal phase to cognitive impairment. Our iNM approach bypasses early or higher lesioned visual pathways, by training redundant, intact, higher cognitive and visual areas to become stronger and functionally associate with lesioned areas. iNM involves learning to perform a physiological response in a controlled and consistent fashion by targeting individual networks when presented with a visual stimulus that serves as both a cue and a neuromodulation interface (**Figure 1A**). This iNM approach is tailored to each participant’s unique anatomical and functional spatial extents and magnitude of blood-oxygen-level-dependent (BOLD; ratio of oxygenated to deoxygenated hemoglobin) signal by extracting and reinforcing maximum or minimum time-resolved signal intensity segment response thresholds under control iNM conditions (**Figure 1C**). By guiding and regulating modulation of the BOLD signal’s spatial extents and magnitude, iNM eliminates the need to rely solely on an individual’s ability to successfully self-regulate the activity of areas that control specific physiologic responses to be neuromodulated.

To determine the mechanisms of visuospatial processing and successfully strengthen the VP network, it was important to develop a task that engages working memory, and selective, divided, attention networks. We thus developed a motion direction discrimination task that engaged visual imagery (VI) in the central vision and VP in the peripheral vision. This task (**Figure SI-2**) required participants to assess: 1) up or down direction of the external stimulus’ motion presented at strong or weak coherence levels in the peripheral vision; and 2) engagement of the internally-generated direction of motion in the central visual space through VI of the peripheral stimulus. Controlling the interplay between peripheral and central vision requires oculomotor planning along with divided attentional processes that regulate error awareness between the constantly adapting VP and VI systems. The rationale to modulate selective attention and working memory networks is based on molecular studies that identified visual system mechanisms while undergoing neuromodulation. The resulting visual spatiotemporal processes that produce functional gains are bolstered by selective attention (also referred as vigilance) and visual perceptual learning (Azimi et al., 2020; Harris & Thiele, 2011; Yousif et al., 2016), as well as longer-term changes in structure and function, such as gene transcription (Gomez et al., 2019; Gu, 2002; Patton et al., 2019; Roelfsema & Holtmaat, 2018).

This study had two primary goals. Because the iNM interface (Figure SI-2) presents the mean intensity and extent of BOLD signal up- or down-regulation in an individual’s unique MT and MST areas (**Figure SI-9**) in the central visual space and the stimulus direction of motion in the peripheral visual space, our first goal was to encode iNM-induced cortical learning mechanisms that connect and control VP and VI. We accomplished this by focusing on attention and memory mechanisms that discriminate direction relative to full and subthreshold motion coherence magnitudes. Our second goal was to decode (Cox & Savoy, 2003) these spatiotemporal system-mechanisms from brain activity during normal (no iNM) and induced (with iNM) learning conditions using pattern recognition (multivoxel pattern analysis), which predicted the probability of upward or downward motion given a brain-state feature vector (Pereira et al., 2009). Thus, to evaluate brain information associated with motion direction and coherence, we trained decoder models to predict the motion direction and coherence of visual stimuli. We used a linear SVM model as our decoder (see Methods for details). To determine whether iNM strengthens the VP and VI networks, we encoded and decoded (i.e., inverse inference, or “brain reading”, Dehaene et al., 1998) physiological and iNM direction discrimination responses at full and subthreshold coherence levels. Neurofeedback studies have provided a causal link between brain activity and physiological responses, as opposed to correlations, making fMRI neurofeedback equivalent to brain stimulation interventions. To elaborate, fMRI neurofeedback can endogenously manipulate brain activity as an independent variable, and it can thus, alter the brain network (Birbaumer et al., 2013; Sitaram et al., 2017; Sulzer et al., 2013; Weiskopf, 2012; T.D. Papageorgiou et al., 2014; T.D. Papageorgiou et al., 2013; T.D. Papageorgiou et al., 2009). Using this analysis. we obtained insights into the *causal relationships* of iNM-induced mechanisms by: 1) quantifying how the brain encodes spatiotemporal features of stimulus direction and coherence to networks; and 2) decoding brain activity by reconstructing coherence and network directions to predict accuracy using control and iNM conditions. Several neurofeedback studies use MVPA or decoded brain responses to define causal brain mechanisms (Haxby et al., 2001; Haynes & Rees, 2005; LaConte et al., 2007; T.D. Papageorgiou et al., 2009, 2013; Shibata et al., 2011; Sitaram et al., 2017).

*Our long-term goal is to use iNM in the clinic to improve visuospatial processing due to low vision (peripheral vision, central vision, or visual acuity deficits) by strengthening VP and VI networks in cortically blind patients or in patients with subjective cognitive impairment (SCI), a prodromal phase to mild cognitive impairment. Currently, there is no effective neurorehabilitative treatment for cortical blindness and no treatment to decelerate the rate of progression for SCI*.

Cortical blindness refers to visual loss due to lesions along retrochiasmal pathways originating below the optic chiasm to the visual cortex. Such lesions produce visual field deficits, including hemianopic or quadrantanopic scotomas (Bruce et al., 2006; Wall, 2021; Zhang et al., 2006) that impede the ability to detect objects within the central visual space (de Jong & Warmink, 2003). Etiological factors of cortical blindness most commonly include stroke (posterior and middle cerebral artery infarct), with an annual incidence of 1 million in the US, of which 27-57% present with retrochiasmal lesions (Gilhotra et al., 2002; Murphy & Corbett, 2009; Pollock et al., 2011). Other factors include head injuries in civilian and military populations, and embolism during surgery (Bruce et al., 2006; G. Poggi, 2000; McKenna et al., 2006; Tierney, 1988). Because the etiology of retrochiasmal lesions is multifactorial, the visual field deficits caused by cortical lesions post-chiasm are more prevalent than those caused by retinal pathology (Dutton, 2003; Zeki et al., 1991). Lesions can be located anywhere along retrochiasmal pathways with varying prevalence depending on the etiology: 12.5% in the primary visual cortex, 23-33% in the optic radiation, 10-19% below the optic chiasm, and 53.6% in various retrochiasmal areas, the bulk of which reside in the occipital lobe.(Bruce et al., 2006; Zhang et al., 2006) Cortical blindness has devastating effects on patient quality of life and productivity, as well as on the economy (Chen et al., 2009; Choi et al., 2022; Dagnelie, 2013; Gall et al., 2010; Jones & Shinton, 2006; E. Papageorgiou et al., 2007). Unfortunately, no current approaches effectively restore vision in these patients (Sabel et al., 2018). Behavioral attempts to restore vision, such as compensatory eye movements, and visual restoration training do not offer practical benefits to patients’ vision (Cowey, 2010; Das & Huxlin, 2010; Horton, 2005a, 2005b; Kasten et al., 2007; Pambakian & Kennard, 1997; Reinhard et al., 2005; Sabel, 2006; Taub et al., 2014; Weiskrantz, 2004; Witte, 1998; Zihl, 1981). In a review of 13 studies investigating treatments for visual field defects, only six compared treatment with a control and of those, only four (67%) showed minimal improvement in reading but not in other tasks of daily living, and there was no reduction in scotoma size (Pollock et al., 2011). Thus, there is no effective treatment to restore cortical blindness deficits.

Subjective cognitive impairment (SCI; (Jessen et al., 2020)), a prodromal phase of mild cognitive impairment (MCI) leading to Alzheimer’s disease, is a state of cognitive decline without objective cognitive deficits on neuropsychological testing or of any other neurological or psychiatric diagnosis (Cheng et al., 2017; Hill et al., 2016; Killeen et al., 2023; Pedro et al., 2016; Wang et al., 2020; Zheng et al., 2018). SCI patients may benefit from visuospatial perception rehabilitation as visuospatial deficits can precede memory impairment in very early phases of cognitive impairment (Zheng et al., 2018). SCI affects 11.2% of adults aged >45 years, while 50.6% of this group also report functional limitations. It is important to decelerate SCI symptoms early, as they are associated with a 1.5-3 fold higher risk of developing MCI or dementia (Mendonça et al., 2016; Ng et al., 2016). SCI diagnostic work-up includes low vision impairment questions, defined as peripheral, and central visual fields deficits. The Behavioral Risk Factor Surveillance System Survey reported that 18% of adults aged 45 years and older, who stated self-reported vision impairment also reported SCD-related functional limitations compared with only 4% of those without vision impairment (Behavioral Risk Factor Surveillance System, 2015). VP deficits that prevent scanning or navigating the environment, such as reading, catching a ball, walking, driving (Okrent Smolar et al., 2023) or, switching between visual tasks, can be due to attention and memory deficits (Saydah et al., 2019; Shang et al., 2021; Vuong & Hedges, 2022; Zheng et al., 2018). Early use of iNM to target the VP network in SCI has the potential to: 1) Diagnose visuospatial impairment, since standard neuropsychological testing is not sensitive enough to identify these deficits; and 2) Decrease the rate of visuospatial deterioration, impacted by a lack of interventions that decelerate functional cortical visual changes.

Because safe and non-invasive new treatments are needed for patient populations with visuospatial and low vision deficits, this study was designed to demonstrate the feasibility of incorporating iNM to neuromodulate VP and VI networks that regulate direction discrimination, as VI has been linked to selective attention and working memory (Albers et al., 2013; Tong, 2013). Using existing knowledge of VP learning and its molecular mechanisms, our experimental design engaged these neural networks with the future goal of applying this intervention clinically. VP is known to be modulated in a bottom-up mechanism, and VI in a top-down fashion (Goebel et al., 1998; Mechelli, 2004; O’Craven & Kanwisher, 2000; Senden et al., 2019). We hypothesized that using a task that activates VP and VI networks through peripheral and central vision, respectively, would strengthen early visual areas that control VP, as well as higher visual, attention, and working memory areas that participate in VI by reinforcing selective exteroceptive and interoceptive attention (Farb et al., 2013), and motor planning networks. We found that the VP-VI iNM interface strengthens motor planning and selective exteroceptive-interoceptive attention by increasing the signal magnitude in early and higher visual areas, as well as parietal and frontal areas. We also found that inverse inference using linear SVMs generated overall higher classification accuracies under iNM across the entire brain and as a function of direction, coherence, and networks than in the control.

## Materials and Methods

### Subjects

Eight healthy, right-handed volunteers (4 males, 4 females, age range = 25-31) were recruited into this 3-day study after obtaining informed consent according to the Baylor College of Medicine Institutional Research Board. Exclusion criteria included prior and current medical or psychiatric diagnoses, intake of any medications, and general contraindications against MRI examinations. Participants had normal or corrected-to-normal visual acuity with MRI-compatible glasses. At the end of each study day, participants were compensated for their time.

### MRI and fMRI Pulse Sequence Parameters

Structural and functional brain imaging was performed at the Core for Advanced Magnetic Resonance Imaging, at Baylor College of Medicine, Houston, Texas using a 3.0 T Siemens Prisma (Siemens, Erlangen, Germany). We used a 20-channel head/neck receiver-array coil to acquire images. A T1-weighted 3D magnetization-prepared, gradient-echo (MPRAGE) sequence acquired 192 high-resolution axial slices [field-of-view (FOV) = 245 x 245 mm^2^; base resolution = 256 x 256; repetition time (TR) = 1,200 ms; echo time (TE) = 2.66 ms; flip angle (FA) = 12°]. Functional data consisted of 33 interleaved axial slices acquired using an Echo Planar Imaging (EPI) sequence (FOV = 200 x 200mm^2^, voxel size = x 3.1 x 3.0 mm, TR = 2000ms; flip angle = 90°, number of volumes = 244).

### Real-time fMRI Neuromodulation Acquisition

Turbo-BrainVoyager (TBV; 2.0; Brain Innovation, Maastricht, The Netherlands) software was used to perform the following five pre-processing computations on EPI images acquired at every time repetition (TR): 1) 3D motion correction; 2) incremental linear detrending to remove BOLD signal drifts; 3) statistical brain map displays generated from a general linear model (GLM) along with beta weights (BOLD signal intensity values) for each condition; 4) extraction of average BOLD signal intensity values from individualized networks acquired on Day 1 scans (see *Data Analysis;* ***Figure SI-9***); and 5) presentation of the network average BOLD signal intensity via the neuromodulation interface (Figure 1A). The ***i***RTfMRIN ***<u>N</u>***euro***<u>M</u>***odulation (***<u>iNM</u>***) interface steps are summarized in Figure 1. To increase the signal-to-noise ratio (SNR), we used an exponential moving average (EMA; Ismailov et al., 2018; Cai et al., amazon web services) algorithm to high-pass filter the ROI BOLD average and suppress low-frequency noise components such as scanner drifts and physiological noise effects (e.g. heart rate and respiration). The EMA output was then low-pass filtered via a Kalman filter to eliminate high frequency noise (Haykin SS. Kalman filtering and neural networks. Wiley Online Library; 2001).

#### Task Design

A random dot kinematogram (RDK) was presented to the lower right quadrant of each subject’s right visual field, while they were asked to fixate on a dot in the middle of the screen (**Figure SI-2a, SI-2b - video**). The RDK displayed upward or downward motion at either fully coherent or subthreshold levels. Four levels of coherent motion were chosen for this study; 100%, 84%, 66%, and 33%. Here we focus on the fully and subthreshold coherence levels represented as C100 and C33 throughout this paper. Using their central vision to fixate on a dot in the middle of the screen, participants were asked to track the direction of RDK motion, which was presented in the lower quadrant of their right visual field, through their peripheral vision as it alternated between direction versus random motion. In the control and neuromodulation conditions, participants were asked to superimpose the upward or downward direction of motion centrally via visual imagery, while direction of motion was tracked via their peripheral vision. In the iNM condition, the central dot served as the neuromodulation interface, i.e., when the central dot filled with red color, it corresponded to successful visual imagery of upward or downward direction of motion. Direction of motion was interleaved with blocks of random motion, during which subjects were asked to rest by disengaging from superimposing imagery of direction of motion while continuing to fixate on the central dot.

#### Study Structure

Our study included two sessions; each consisting of ten functional (echo planar imaging; EPI) scan, which included five control-no iNM scans that alternated with five neuromodulation (iNM) scans. Each EPI scan included eight continuous periods each lasting 8 minutes and 12 seconds. Within each period, subjects were cued to imagine motion perception as either up or down depending on the RDK session displaying one of the four coherence levels. Each coherent motion block lasted 20 secs and was interleaved with a baseline-random motion block (10 secs). The direction of coherent motion blocks was randomly counterbalanced across runs, following three rules: 1) each coherent motion block occurred twice during each period; 2) a coherent motion block was never followed by the same coherence level; and 3) each run consisted of a unique block order.

#### Neuromodulation Paradigm

Neuromodulation was determined by color and extent of a circle that was filled, representing the magnitude and extent of each subject’s targeted network. The neuromodulation signal was calculated by comparing the <u>p</u>ercent BOLD <u>s</u>ignal <u>c</u>hange (PSC) generated during each control run for each coherence block with the rest block that preceded it. The BOLD PSC change was calculated from each participant’s individualized areas every 2 seconds as follows: *BOLD PSCi(j) = 100% * [ROIs BOLD during Up OR Down direction selectivity i(j) –ROIs BOLD during tongue at rest i(j)]tongue rest i(j)* where *i* represents coherence level (C100; C84; C66; C33), *j* represents the time interval (2 secs) used to compute the BOLD PSC of coherence level *I* . The neuromodulation presented at each TR was computed by comparing the current PSC value with a PSC reference range that included seven bins of 25% BOLD increase or decrease: -100%; - 75%; -50%; -25%; 0; 25%; 50%; 75%; 100%. During the iNM run following each control run, if the PSC at a given time point was within the reference range or higher than the maximum value, the circle was filled with red, representing upregulation of the targeted ROI BOLD signal that controlled visual perception and imagery. If the PSC was lower than the minimal value, the circle was filled with blue, representing downregulation of the targeted ROI BOLD signal that controlled visual perception and imagery.

#### Analysis

Acquired anatomical and functional images were processed offline using AFNI (**A**nalysis of **F**unctional **N**euro**I**mages; Cox, 1996; http://afni.nimh.nih.gov/afni). Anatomical data was spatially transformed to Talairach space using the TT_N27 atlas. Functional data was preprocessed to reduce artifacts and increase SNR. Our signal pre-processing protocol included: 1) removal of outliers (head motion, physiological artifacts; 3dDespike0 from the time series; 2) slice-time correction (3dTShift); 3) transformation of our oblique-acquired functional dataset to a cardinal dataset (3dWarp - oblique2card); 4) motion correction by registering each functional dataset to the first volume of the first functional run, using a 3D-rigid-body transformation; 5) spatial smoothing with a 6 mm-FWHM Gaussian kernel filter; 6) co-registration of functional data with individual T1-weighted 3D-structural data; and 7) scaled to have a mean BOLD signal intensity of one-hundred.

Statistical analysis was performed on fMRI data using a GLM approach. Five runs for each condition (i.e. control or iNM) were then concatenated to increase the SNR prior to generating parametric brain maps across different coherence levels for each patient. A second-order polynomial was used to model slow baseline shift. Model parameters were estimated using the 3dREMLfit program in AFNI, which uses an ARMA(1,1) model to estimate the correlation structure in noise. The GLM included: 1) four regressors, one for each of the four coherence levels (C100, C84, C66, C33); 2) six covariate vectors that controlled for head motion; 3) regression of white matter and cerebrospinal fluid means to increase SNR, since activity in these areas represents noise; and 4) baseline-random motion blocks. The four regressors corresponded to each coherence level and were convolved with a gamma variate function, a canonical hemodynamic response function. Group analysis was then performed using 3dMEMA, a mixed effects meta-analysis program that models within and across subject variability. After generating a network of regions for VP-VI control compared to rest-baseline (voxelwise p-value=.05), cluster analysis was performed to identify highly significant ROIs (FWER<0.01) using a cluster-size of at least 10 voxels (3dClustSim, AFNI).

The remaining analysis was performed in Python. We focus here, in the analysis for coherence levels C100 and C33. Significant ROIs for each coherence level and motion direction were analyzed and certain ROI clusters that represented dataset noise were removed. Anatomical masks as defined in the AFNI TT_Daemon atlas were created and resampled for alignment with participants’ functional scans. These functional masks created after Day 1 from the localizer scans were then used to extract statistically significant ROIs that corresponded to the middle temporal (MT), and middle superior temporal (MST) cortices. Functional data was detrended using a second-order polynomial before being scaled to a mean BOLD signal intensity of 100. Time series BOLD responses were extracted across the entire brain for analysis to generate significant networks formed from the control and iNM conditions. Extracted time series signals for specific brain regions were used to conduct the remaining analyses.

#### Networks

In this study, we defined four distinct networks: Visual Perception, Visual Imagery, Motor Planning, and Intero-Extero-ceptive Attention. The individual brain regions contained within each network are listed under figure **SI-1**.

### Support Vector Machine Classification

A linear support vector machine (SVM) was used to decode BOLD spatial patterns of each coherence level (i.e. C100 and C33) generated by the five neuromodulation and five control conditions. Each functional run included 120 3-dimensional participant fMRI scans representing the BOLD signal at a particular TR (120 TRs total). Within each run, the selected coherence level was labeled 1 and the random motion block following the selected coherence level was labeled 0. If the selected coherent motion block was at the end of the run, the preceding random motion block was used as the baseline and labeled 0. The SVM was trained to distinguish between these two classes for each coherence level. We used a regularization parameter, c, of 0.0001 based on a hyperparameter search and applied a 5-fold cross-validation procedure to each neuromodulation condition. For each subject and each coherence level, we trained the SVM model on the concatenated data from four runs of the corresponding condition (i.e. control or iNM) and tested its performance using area under the receiver-operator curve (AUROC) on the remaining run. This was repeated in a cross-validation approach and the AUROC’s were averaged to achieve a single AUROC for each subject with a coherence level. Differences in classification accuracies were compared across coherence level conditions and a paired t-test was run using the AUROC values for each subject. SVM was also used to decode BOLD spatial patterns within predefined networks (VP, VI, MP, SEIA) for each coherence level.

### Area Under the Curve

To compute the BOLD PSC for each coherence level and direction of motion as a function of time, ROIs were categorized across VP, VI, MP, and SEIA networks. BOLD PSC time series were generated by averaging: 1) the time series of voxels within an ROI; and 2) over signal time-segments from blocks of the same coherence level collected from runs of the same direction. Each signal segment started at the onset of a coherent motion block and ended at the last time point of the following random motion block with the exception of the last block of the scan, which used the random motion block preceding the coherent motion block, since the scan ended with a coherent motion block. Each subject had five control and five iNM runs with each run entailing two blocks of VP-VI at each coherence level. The PSC time series for each coherence level, each condition, and each subject was averaged over ten BOLD signal segments. The area under the PSC curve of each condition (control or iNM) was calculated using Simpson’s Rule:

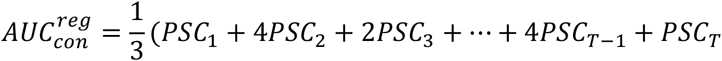

Where PSC_t_ is the PSC at the corresponding time t.

The BOLD PSC was then calculated using the following formula:

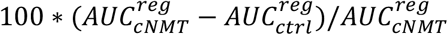

### D’ Sensitivity Index

To measure the strength of iNM across time and rank regions, we used a variant of Cohen’s d known as the D’ sensitivity index. This index quantifies the separation between the control and iNM distributions means reslative to their standard deviations. Assuming that the fMRI data set has *T* time points, *B* blocks, *P* subjects, and *N* ROIs, each ROI was characterized by a 3-D matrix of dimensions *T* × *B* × *P*. Each subject was then characterized by a 2-D matrix of dimensions *T* × *B*. For a fixed time point *t*, the distributions of control and iNM blocks were 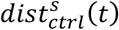 and 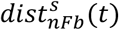 respectively. D’ for the subject s at time point *t* is:

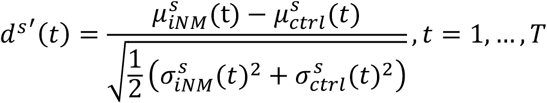

Where 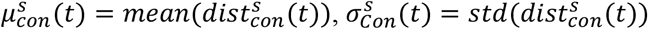.

The sensitivity index between the two conditions at time point *t* and for the fixed region is the average sensitivity index across the subjects:

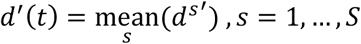

## Results

### iNM Improves Classification Performance Over Control-no iNM Condition Using Linear Support Vector Machine Analysis

Using a linear SVM, we aimed to predict the coherence level presented to participants using BOLD signal intensity. We found that a linear SVM could significantly improve classification performance under neuromodulation as shown by the median AUROC under all coherence levels and motion directions. The median AUROC for each coherence level and motion direction was: 1) .68 AUROC iNM vs .52 AUROC control for **C100 up** (p-value: 0.001); 2) .71 AUROC iNM vs .53 AUROC control for **C100 down (**p-value: 0.004); 3) 64 AUROC iNM vs .54 AUROC control (p-value: 0.045) for **C33 up**; and 3) .63 AUROC iNM vs .55 AUROC control (p-value: 0.048) for **C33 down (Table S5a)**.

When we tested the effect of including VP, VI, MP, and SEIA networks within the classification scheme, the classification performance of two networks in the C100 up condition was higher under neuromodulation than in the control condition: MP network (.63 AUROC iNM vs .55 AUROC control; p-value: 0.05) and SEIA network (.61 AUROC iNM vs .53 AUROC control; p-value: 0.004). The remaining two VP, VI C100 up networks were trending toward significance but did not achieve it: VP (.58 AUROC iNM vs .55 AUROC control; p-value: 0.06) and VI (.59 AUROC iNM vs .56 AUROC control; p-value: 0.08). In the C33 up condition, three networks achieved significance: VI (.59 AUROC iNM vs .51 AUROC control; p-value: 0.07), MP (.63 AUROC iNM vs .50 AUROC control; p-value: 0.02), and (.65 AUROC iNM vs .52 AUROC control; p-value: 0.004). The VP network for C33 was trending toward significance but did not achieve it (.59 AUROC iNM vs .51 AUROC control; p-value: 0.07). All C100 down networks were statistically significant: VP (.61 AUROC iNM vs .50 AUROC control; p-value: 0.006), VI (.64 AUROC iNM vs .54 AUROC control; p-value: 0.02), MP (.68 AUROC iNM vs .49 AUROC control; p-value: 0.0007), and SEIA (.72 AUROC iNM vs .49 AUROC control; p-value: 0.0008). No networks in the C33 down condition achieved or trended toward significance: VP (.56 AUROC iNM vs .56 AUROC control; p-value: 0.32), VI (.62 AUROC iNM vs .54 AUROC control; p-value: 0.29), MP (.57 AUROC iNM vs .56 AUROC control; p-value: 0.32), and SEIA (.61 AUROC iNM vs .60 AUROC control; p-value: 0.86).

The percent change in AUC signal intensity illustrates the effect of neuromodulation on signal intensity over the course of each block in the scan (**Figure 3**). The effect of neuromodulation was most varied on VP and VI networks. The VP network was significantly lower at coherence level C33, with a -1362% change in AUC signal intensity for the up direction and -124% for the down direction. The VI network was also lower at coherence level C33, with a 4% decrease in the up direction and a 15% decrease in the down direction. Interestingly, at coherence level C100, no significant VP network activity was seen in either direction in the control condition, with significant VP network activity only seen during neuromodulation. At coherence level C100 down direction, the VI network in the iNM condition showed a 19% increase in AUC signal intensity. while in the up direction, AUC signal intensity was 47% higher compared to the control. The MP network was upregulated at all coherence levels and directions in the iNM condition compared to the control with AUC signal intensity ranges from 26-59%. iNM SEIA networks also increased with AUC signal intensity ranges from 48-76%. Detailed analyses of each ROI for each network across all coherence levels and directions are shown in the Supplemental Information (**Figures SI-3 – SI7**).

**Figure 2.**
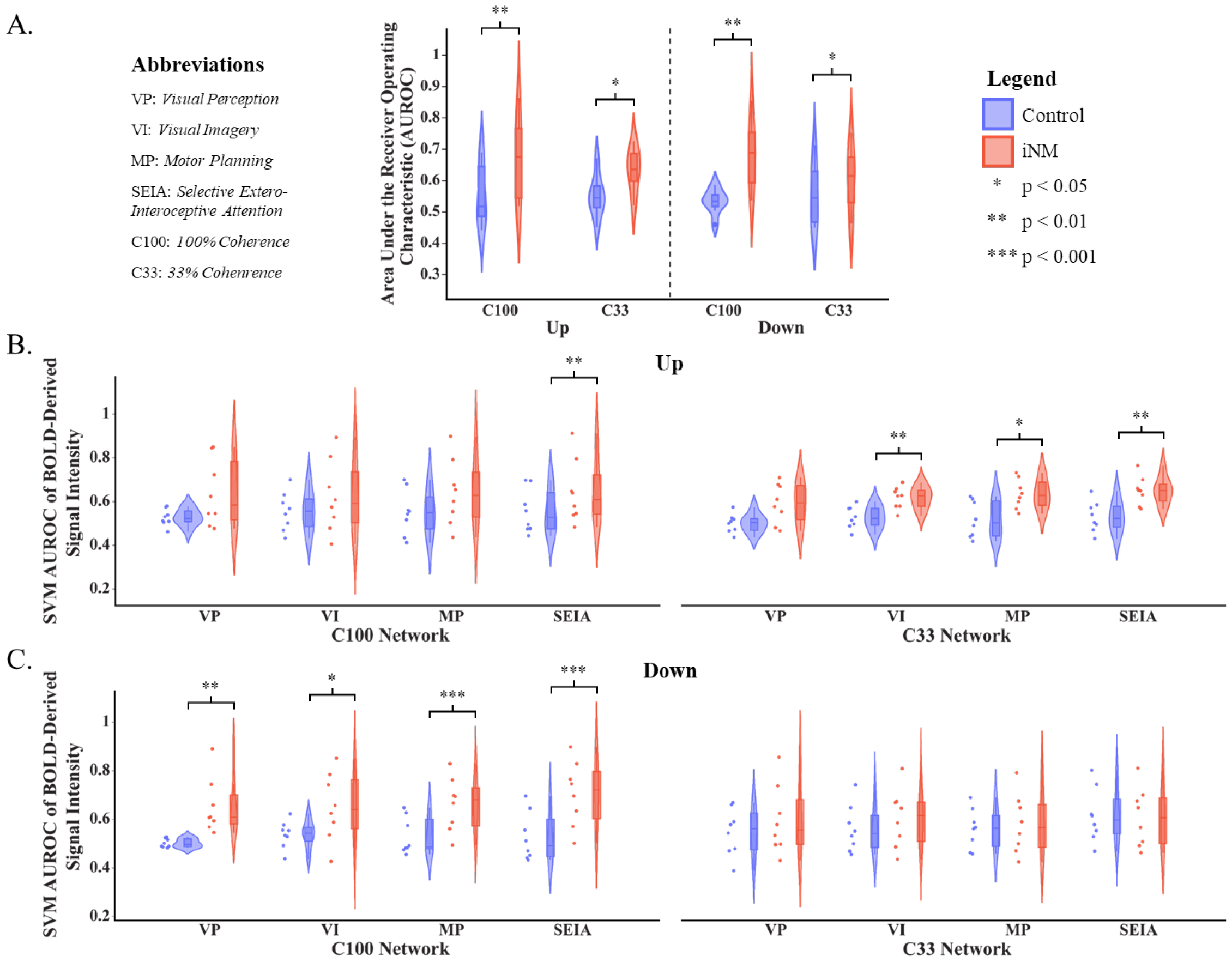
Increased classification performance in iNM condition over control-no iNM condition for all coherence levels and networks using a linear support vector machine. **A**. Violin plots of SVM-generated areas under the receiver-operator curve (AUC-ROC, y-axis) performance values across all subjects relative to coherence level (x-axis) and training-testing data type permutations (color-coded). On each box, the central line, and lower and upper edges represent the median (Q2), 25th (Q1), and 75th (Q3) percentiles, respectively. The dots represent outliers (bigger than Q3 by 1.5*(Q3-Q1) or smaller than Q1 by 1.5*(Q3-Q1)). Whiskers extend to the most extreme non-outlier datapoints. Black lines and asterisks indicate significant differences by paired-samples Wilcoxon sign tests. **B**. Violin plots of SVM-generated AUC-ROC performance values across all subjects relative to C100 and C33 coherence levels in up direction cortical networks. **C**. Violin plots of SVM-generated AUC-ROC performance values across all subjects relative to cortical networks in C100 and C33 down direction coherence levels.

**Figure 3.**
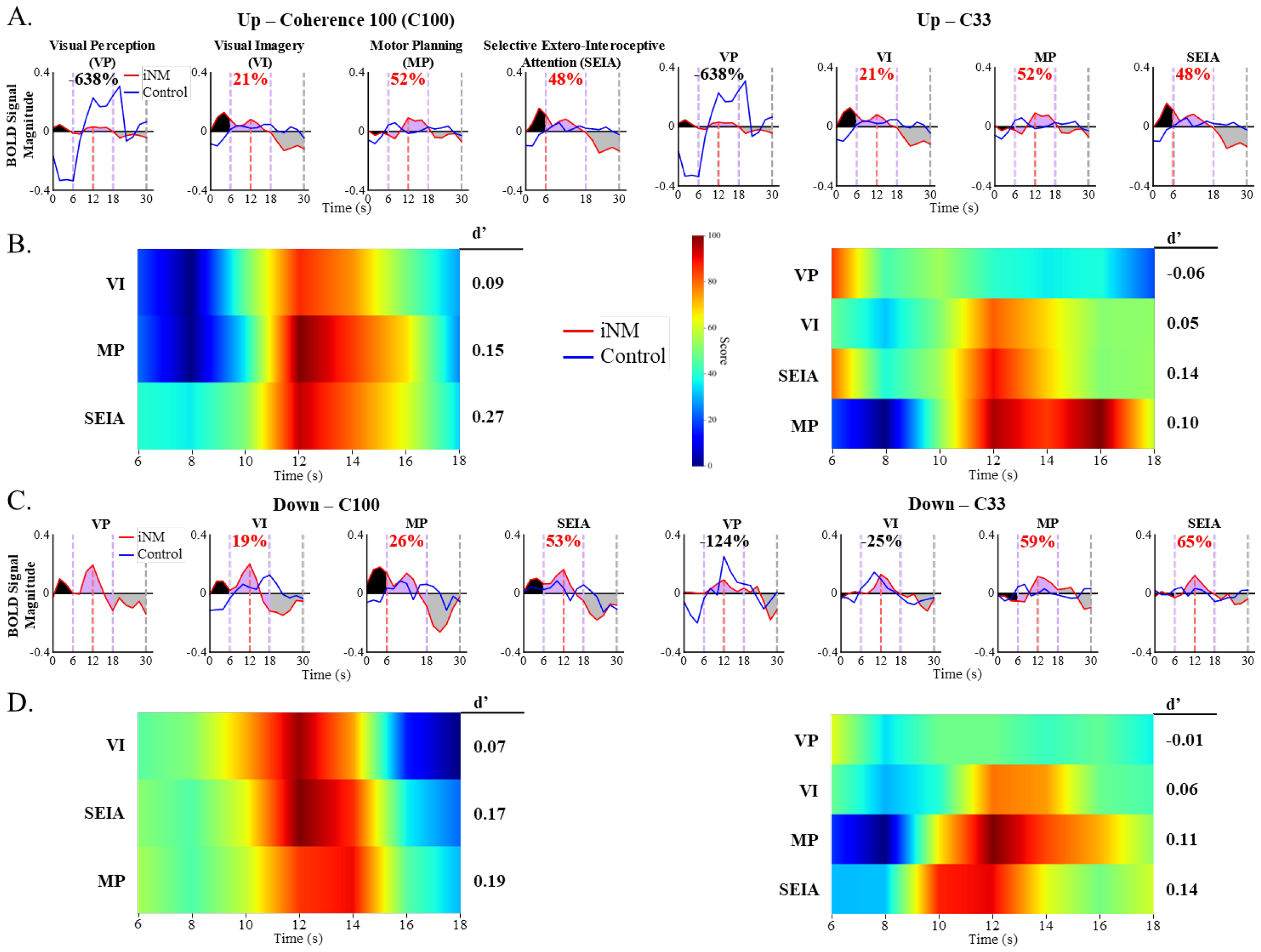
iNM increases mean AUC signal intensity and d’ sensitivity indices compared to control conditions across networks, motion directions and coherence levels. A. BOLD signal intensity of both iNM and control conditions are plotted as a function of time for all networks and coherence levels in the up-motion direction. Each plot contains three sections: 1) a hemodynamic lag block (0-6 seconds), 2) a coherent motion block, which was truncated to 18 seconds due to participant fatigue in the task, as the signal decreased across all ROIs (6-18 seconds), and 3) a baseline-random motion block (18-30 seconds). The area under the BOLD signal intensity curve of the iNM condition is colored black to indicate the hemodynamic lag; purple for coherent motion; and gray for baseline-random motion blocks. Percent change in area under the curve between iNM and control conditions for the coherent motion block is in red. B. The d’ sensitivity index plotted as a function of time in the coherent motion block for all networks and coherences in the up-motion direction. The d’ sensitivity value for the entire period is shown to the right of each heatmap. C. BOLD signal intensity of iNM and control conditions plotted as a function of time for all networks and coherence levels in the down-motion direction. D. The d’ sensitivity index plotted as a function of time in the coherent motion block for all networks and coherences in the down-motion direction.

The d’ sensitivity index was also computed for each experimental time point in the form of heatmaps. The d’ sensitivity index measures the separation between the means of the control and iNM distributions relative to their standard deviations. For the C100 up condition, all three networks (VI, MP, and SEIA) had similar d’ sensitivity index profiles, with greater activation in the control condition after 10 seconds, at which time activation under the neuromodulation condition increased above the neuromodulation condition. For the C33 up condition, the VP network peaked first, followed by peak activation of VI, and SEIA around 12 seconds, and finally MP at 16 seconds. Activation of all networks in the C100 down condition peaked at 12 seconds; however, activation of the VI network started earlier than in the SEIA and MP networks and also began to ramp down earlier. Activation of SEIA and MP lasted longer (until 15 seconds). Finally, in the C33 down condition, VP was activated first followed by the SEIA network and simultaneous peak activation of the VI and MP networks. However, activation of the MP network began earlier, near the start of SEIA network activation.

## Discussion

The purpose of designing a visual perception-visual imagery (VP-VI) paradigm and neuromodulation interface was to strengthen early and higher visual perception, as well as selective attention and working memory networks. To determine the spatiotemporal mechanisms of induced learning via iNM, we encoded stimulus data and decoded brain data, to present a causal inference to the spatiotemporal mechanisms. Our long-term goal is to apply this visuospatial iNM approach in two patient populations for which no effective, safe, and non-invasive intervention exists, with the goal to neurorehabilitate low vision deficits, in cortically blind patients with early or higher VP lesions, and to decelerate visuospatial deficits in patients with prodromal SCI prior to presenting with signs of cognitive deterioration.

This VP-VI task engaged: 1) the peripheral visual field, involving visual perception of the direction of randomly moving dots at strong and weak coherences; and 2) the central visual field, involving VI of motion direction and superimposing that imagery with the internally-generated stimulus at central eye fixation where the iNM interface was located (**Figure 1A**). We found that iNM resulted in greater sensitivity index (also referred as d prime, d’) in the regions activated under strong coherence, independent of direction. The BOLD magnitude of the AUC for VP and VI networks across directions at full threshold (100%) coherence was also greater under iNM. However, at weak, subthreshold (33%) levels, the VP AUC decreased in both directions under iNM, while the VI AUC network did not change in the up direction but decreased in the down direction (**Figure 3**). Thus, in response to subthreshold coherence in a field of randomly moving stimuli (dots), the BOLD signal decreased VP AUC for both up- and down-motion directions under iNM. Similar findings were seen in microstimulation studies when low coherence stimuli were not reliably identified, while the strength of the motion signal in a field of randomly moving dots varied during a direction discrimination task (Ditterich et al., 2003; Nichols & Newsome, 2002).Whether or not intensity of the VP and VI networks was modulated as a function of coherence under iNM, both directions across strong and weak coherences were accompanied by increased BOLD magnitude in the motor planning (MP) and selective interoceptive-exteroceptive attention (SEIA) networks.

Visual perception requires communication between ongoing spontaneous activity in visual cortex and activity evoked by a stimulus (Hesselmann et al., 2008). That is, the extent and intensity of visual cortex activity while a stimulus is presented depends on the magnitude of VP (Ress et al., 2000). In this study, we presented the stimulus direction of motion as a function of full (100%) and subthreshold (33%) coherence. When motion coherence was weak, visual cortex activity was weaker under iNM than in the control. Also, when coherence was weak, the magnitudes of the MP and SEIA networks increased under iNM in the up and down directions. Our findings agree with new evidence that motor and somatosensory areas predict information about eye movements in early visual areas (Guitchounts et al., 2020; Leinweber et al., 2017; Mertens et al., 2023). This suggests that the VP network transmits and allocates a signal to MP and SEIA networks to fulfill task demands that control oculomotor activity between the peripheral VP network and the VI performed in central space. Thus, the lack of increased VP under iNM could be due to outflow and higher resource allocation to the MP and SEIA networks to fulfill task demands, rather than to the control.

Activity generated by MP areas seems to represent oculomotor planning and control between tracking the VP stimulus presented in the peripheral space and the internally-generated, VI stimulus at central eye fixation. Common MP areas across directions and coherences included the precentral gyrus and anterior motor cerebellum (represented by the fastigial nucleus), both of which are needed for persistent cortical preparatory activity (Gao et al., 2018). The precentral-primary motor area, anterior cerebellum, basal ganglia, and supramarginal gyrus have been implicated in higher motor cognition and willed generation of visual sensory and motor imagery (Grèzes & Decety, 2001; Hesse et al., 2006).

Neurons in the primary motor cortex, also known as the precentral area, respond to visual stimuli when used to initiate and plan a movement. The basal ganglia play a key role in voluntary saccadic eye movements (Hikosaka et al., 2000) via inhibitory signals that travel from the caudate to the substantia nigra (SN) and from the SN to the superior colliculus. Macaque studies have shown that in the presence of a reinforcement mechanism, the basal ganglia can modify oculomotor control (Sato & Hikosaka, 2002). In our study, iNM provided a reinforcement mechanism, during which the caudate and globus pallidus signal intensity increased when compared to the control.

Anterior cerebellar activity was also enhanced under iNM treatment. It is well established that the cerebellum is instrumental in controlling eye movements and visually-guided motor learning, specifically influencing perception and motor control as it adjusts the acquisition of perception through projections to sensorimotor areas, such as the premotor, supplementary motor area, and precentral and primary somatosensory cortex (Brodal, 1978; Glickstein et al., 1985; J. D. Schmahmann & Pandya, 1989). There are various hypotheses about cerebellar processing as a forward or inverse internal model. A forward model predicts oculomotor position, while an inverse model involves planning appropriate eye movements between peripheral and central visual space according to task demands (Glasauer, 2003). There is no explicit support for either of these hypotheses, but each provides a partial or a synergistic model to interpret the cortical activations. Our results support the view that persistent neural dynamics during MP are maintained by multi-regional neural circuits (Guo et al., 2017), such as the prominence of anterior motor cerebellar computations representing online motor control posed by the task demands (Guo et al., 2017; J. Schmahmann, 1998).

We also found that modulating visual sensory direction and magnitude of coherence inputs interpreted by VP and VI networks, strengthen areas involved in crosstalk between selective exteroceptive and interoceptive attention (SEIA) networks during iNM-induced learning. Crosstalk between these two attention systems controls conscious perception-awareness of one’s errors (Ullsperger et al., 2010). Attention can be driven exogenously or endogenously (Berger et al., 2005; Chong & Blake, 2006), with selective attention divided into focused and divided attention. In this study, exteroceptive attention was driven exogenously via incoming interpretation of peripheral sensory information through the VP network, and interoceptive attention was driven endogenously via internally-generated VI superimposition of the stimulus. Thus, selective attention was divided, as these two systems must constantly control the interplay between selective exteroceptive (VP) and interoceptive (VI) attention. iNM enhanced the intensity of selective attention areas, encompassing communication between: 1) exteroceptive attention via incoming peripheral vision sensory information from external stimuli by activating visual motion perception areas; and 2) interoceptive attention to internally-generated stimuli by superimposing VI of up or down directions at full and subthreshold levels.

Decreased VP magnitude of the BOLD signal when coherence is weak under iNM is driven by various factors. One hypothesis is that interplay between ongoing activity in the early visual cortex and that evoked by the stimulus could trigger a bottleneck as perceptual demands must be allocated to cues displayed by the iNM interface. Previous studies have shown that VP depends on the interaction between visual cortex activity and the nature of the stimulus (Ress et al., 2000). Another, perhaps more plausible, hypothesis is that although iNM decreased the BOLD magnitude of VP, the magnitude of the VI, MP, and SEIA networks increased, suggesting that balanced allocation of resources is required under iNM to successfully complete task demands. It is possible that decreased VP for weak coherence could result from allocation of VP to the networks needed to meet demands of the visual imagery task, which required MP and selective attention processes of externally-presented peripheral visual space and internally-generated stimuli at central visual space. iNM increased the BOLD magnitude of MP (up; 52% and down: 58%) and SEIA (up: 48% and down: 65%) networks, suggesting that to effectively allocate the physiological demands of this complex imagery task requires engagement of working memory to generate an internal visual imagery stimulus along with selective intero-extero-ceptive attention and MP. Thus, similar to previous reports (Brefczynski & DeYoe, 1999; Kastner et al., 1998; Kosslyn et al., 2001; Slotnick et al., 2005; Stokes et al., 2009) constant communication between bottom-up and top-down control networks is required. Recruitment of anterior motor and posterior sensory (J. Schmahmann, 1998) cerebellar regions is also required because successful VP depends on interplay between ongoing spontaneous visual cortex activity with that evoked by a stimulus (Hesselmann et al., 2008).

## Conclusions

**1**. iNM enhances the magnitude of VI BOLD motion direction by increasing the engagement of memory-related signals, such as those of the hippocampus, frontal, intraparietal, and superior parietal areas. Similar findings have been reported in macaque studies (Zaksas et al., 2001; Zaksas & Pasternak, 2006b).

**2**. iNM enhances SEIA modulation (Farb et al., 2013) across motion directions and coherences by engaging posterior cerebellar (pCb) areas (SI-5-7). pCb activity has also been shown to participate in attention and working memory demands (Leggio et al., 2008; Peterburs et al., 2019; Restuccia et al., 2006).

**3**. iNM enhances the ability to perform visual imagery of strong and weak motion information by engaging SEIA and MP networks. These findings are consistent with the dorsal attention network, which includes the pCb, visual motion areas, frontal eye fields, parietal lobule, intraparietal sulcus, and premotor cortex (Buckner et al., 2011).

**4**. When coherence was weak, the VP network BOLD magnitude decreased compared to the control while resource allocation – BOLD intensity – appears to have been distributed to MP and SEIA networks to adaptively monitor oculomotor and attentional demands behavior between peripheral vision, and central vision performing VI, in response to iNM demands.

**5**. Under control conditions, and weak coherence, the brain has greater VP demands (BOLD intensity) when interpreting direction of motion, as early visual areas displayed more activity than under iNM in response to the awareness and perception of motion, while decreased intensity of VI, MP, and SEIA networks suggests that resources for selective attention and working memory are significantly dampened. Similar findings have been observed in macaque studies (Zaksas & Pasternak, 2006b).

## Limitations of present study

Although the number of participants who received iNM was small, we have decoded direction, coherence, and network participation, finding statistically significant differences between iNM and control-no-iNM. Although we did not have an MR-compatible eye-tracker to measure eye movement due to prohibitive cost, we strategically placed the iNM interface at a central location so that participants’ eyes would fixate in the central part of the visual field and only track the motion presented in their peripheral space. If participants were not able to fixate centrally, it would have affected their iNM feedback and researchers would have been aware through evolution of the time series every 2 seconds, which would have eventually plateaued. Although we lack a behavioral measure to determine participants’ direction discrimination, we were able to assess performance using computational models that quantified the encoding networks generated and decoding the prediction of performance. These computational models agree in their findings as a function of direction, coherence, and network.

