## Supplemental Information for "Individualized fMRI neuromodulation enhances visuospatial perception: a guided approach targeted towards the neuro-rehabilitation of cortical blindness and deceleration of subjective cognitive impairment"

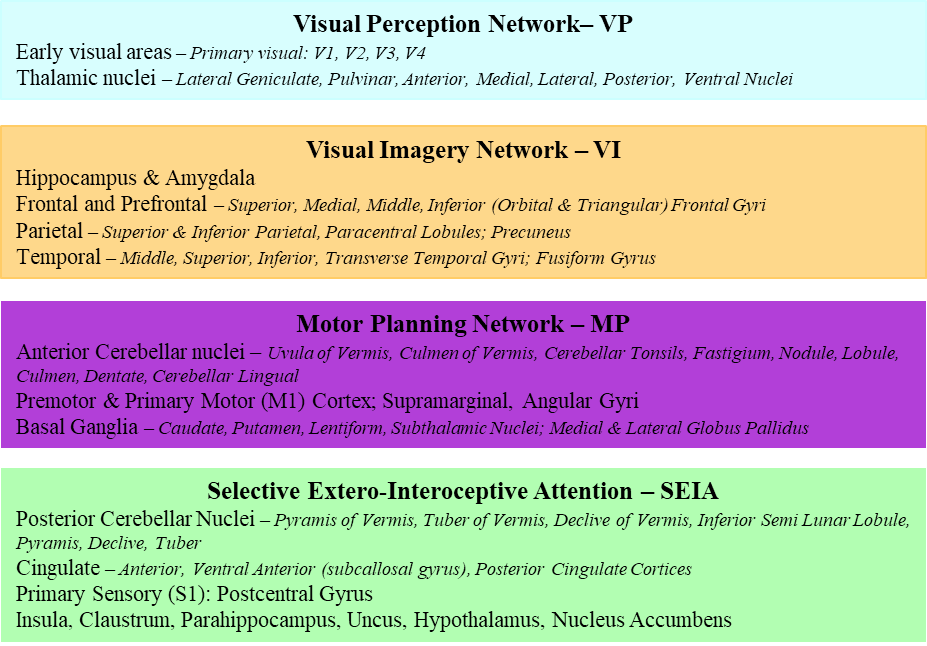
**Supplemental Information – Figures and Tables**

**Figure SI-1.** List of individual regions within each network.

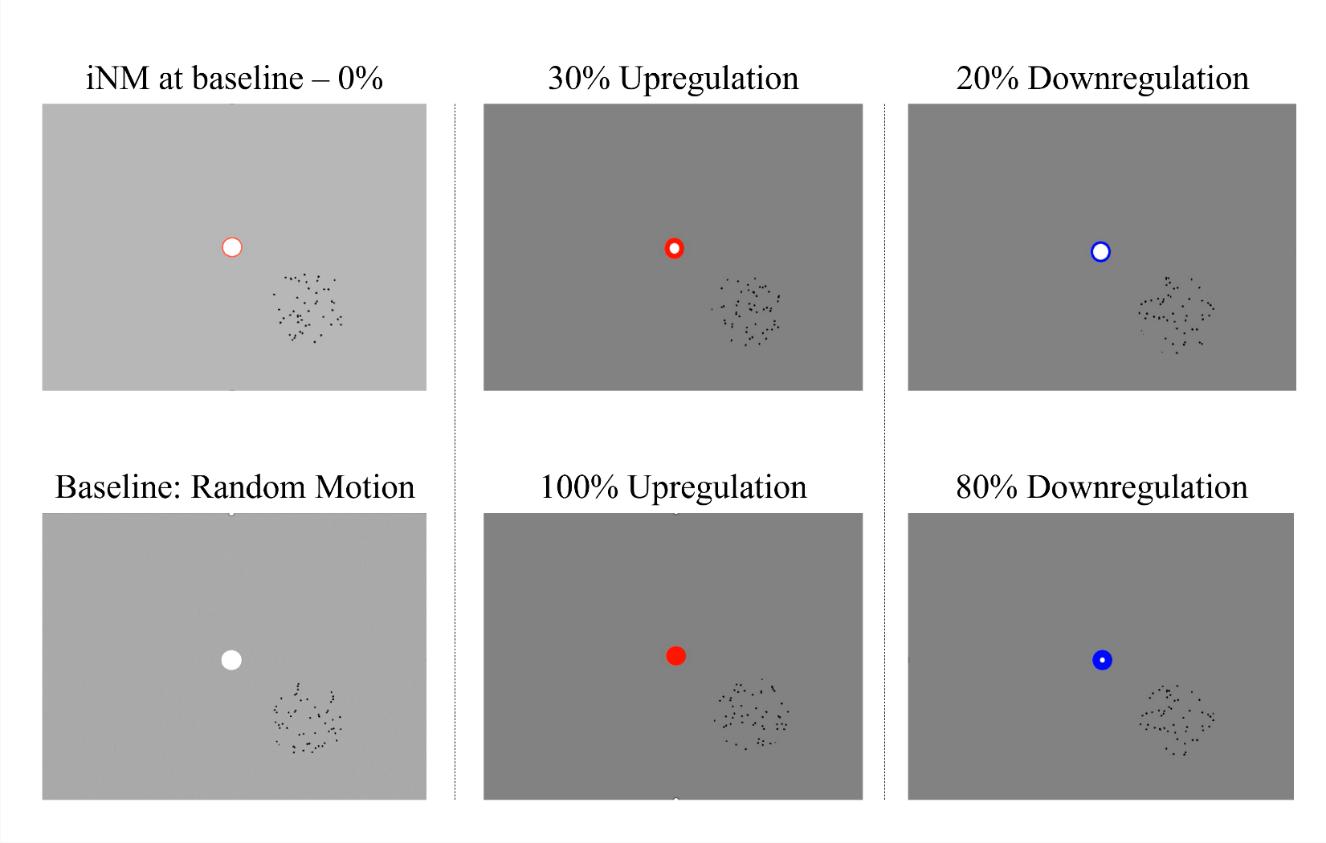
**Figure SI-2a.** Video of iNM interface.

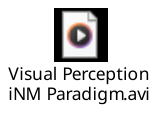

**Figure SI-2b.** Video of iNM neuromodulation interface.

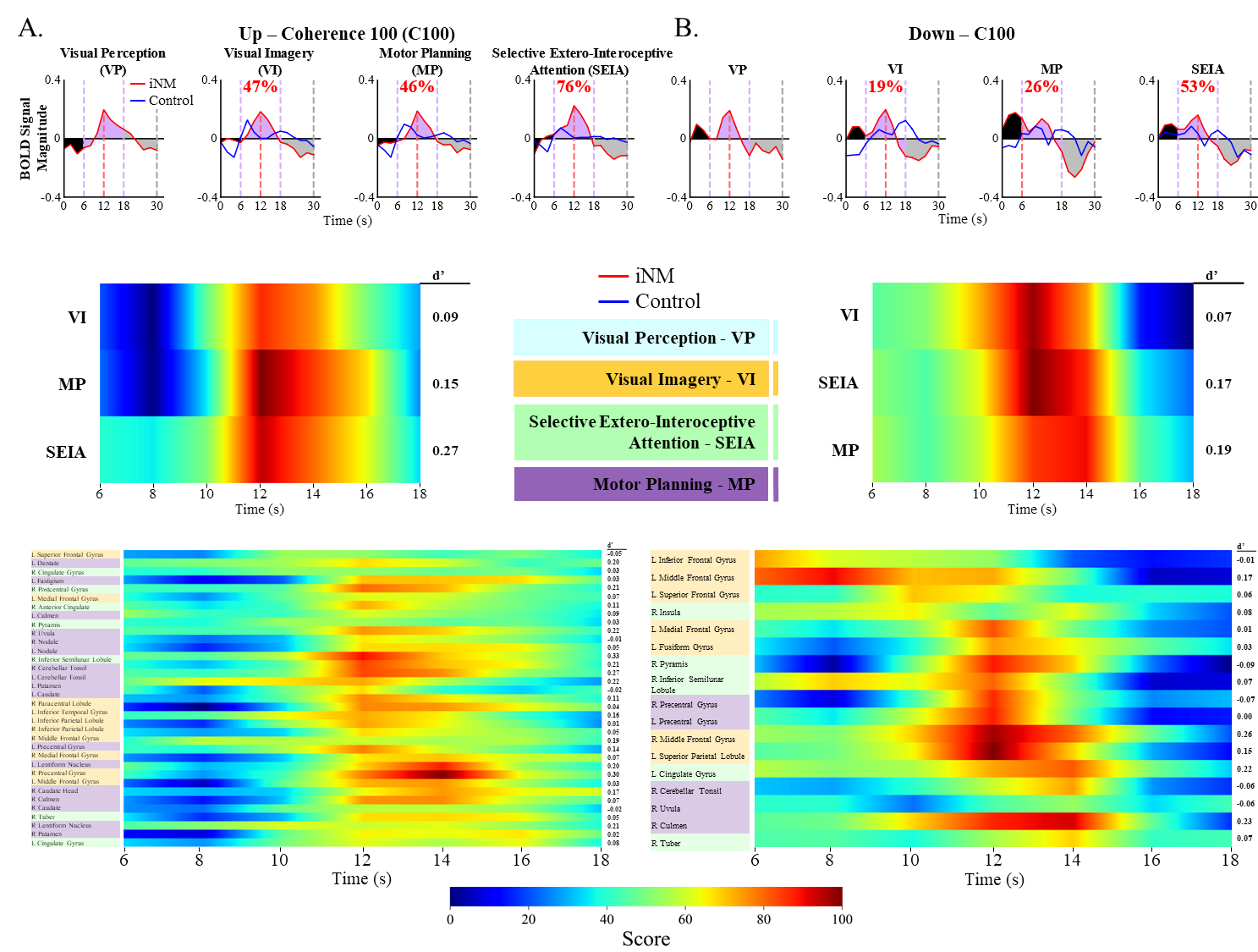
**Figure SI-3.**

A. BOLD signal intensity of iNM and control conditions are plotted as a function of time for all C100 coherence level networks and motion directions. Each plot contains three sections: 1) a hemodynamic lag block (0-6 seconds), 2) a coherent motion block that was truncated to 18 seconds, as the signal intensity decreased across all ROIs (6-18 seconds) most likely due to participant fatigue in the task, and 3) a baseline-random motion block (18-30 seconds). Area under the iNM condition BOLD signal intensity curve was colored (black, hemodynamic lag; purple, coherent motion; or gray, baseline-random motion blocks. Percent change in the area under the curve for the coherent motion block control and iNM conditions is shown in red.

B. The d’ sensitivity index is plotted as a function of time in the coherent motion block for all networks and motion directions in coherence level C100. The d’ sensitivity value for the entire period is shown to the right of each heatmap.

The d’ sensitivity index is plotted as a function of time in the coherent motion block for all individual regions and motion directions in the C100 coherence level. The d’ sensitivity value for the entire period is shown to the right of each heatmap. Color of the individual region name corresponds to the region network.

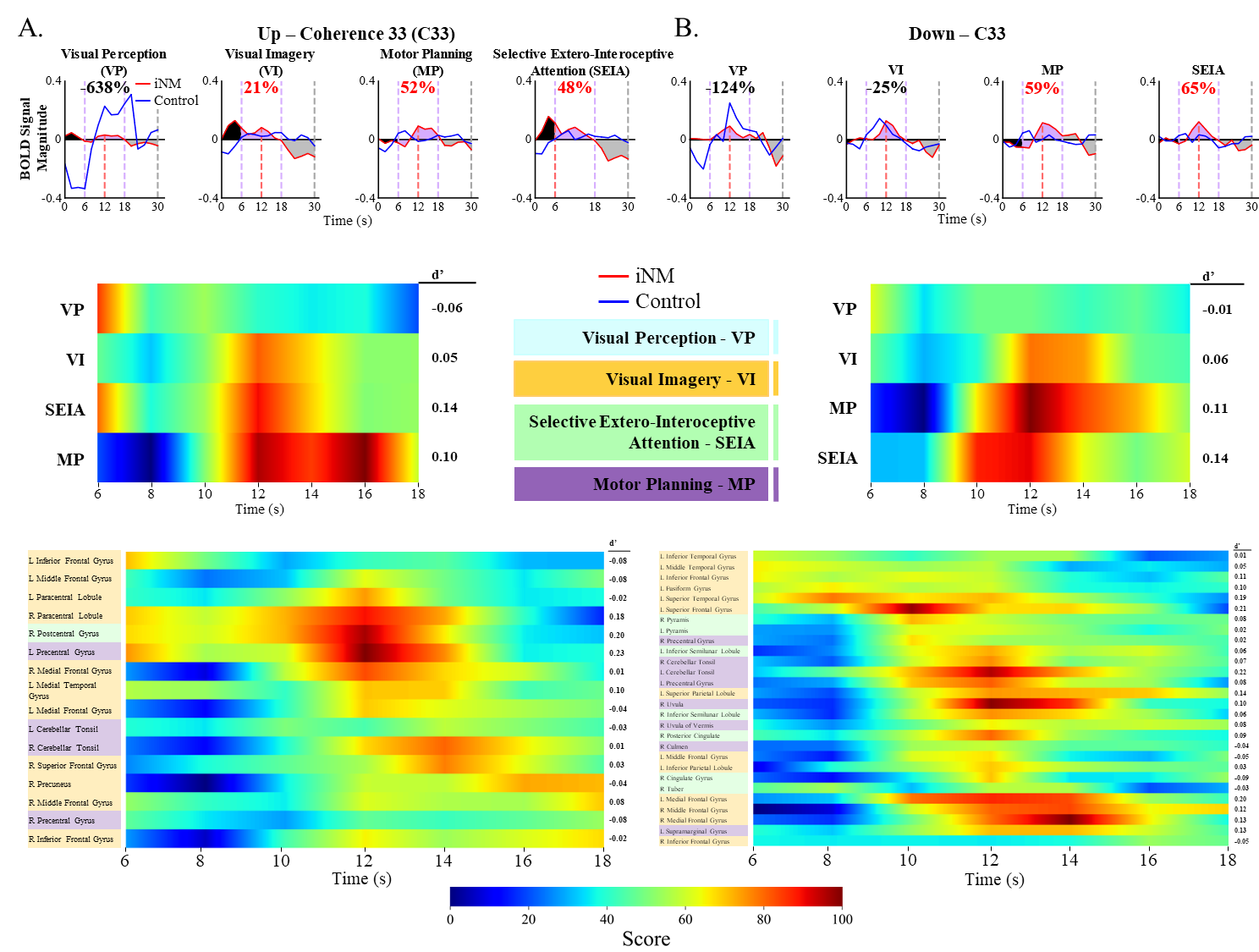
**Figure SI-4.**

A. BOLD signal intensity of iNM (red) and control (blue) conditions are plotted as a function of time for all C33 coherence level networks and motion directions. Each plot contains three sections: 1) a hemodynamic lag block (0-6 seconds), 2; a coherent motion block that was truncated to 18 seconds, as the signal intensity decreased across all ROIs (6-18 seconds) most likely due to participant fatigue in the task; and 3) a baseline-random motion block (18-30 seconds). The area under the iNM condition BOLD signal intensity curve was colored: black, hemodynamic lag; purple, coherent motion; or gray, baseline-random motion blocks. Percent change in area under the curve between the coherent motion block control and iNM conditions is shown in red.

B. The d’ sensitivity index is plotted as a function of time in the coherent motion block for all networks and motion directions in coherence level C33. The d’ sensitivity value for the entire period is shown to the right of each heatmap.

The d’ sensitivity index is plotted as a function of time in the coherent motion block for all individual regions and motion directions in coherence level C33. The d’ sensitivity value for the entire period is shown to the right of each heatmap. Color of the individual region names corresponds to the region network.

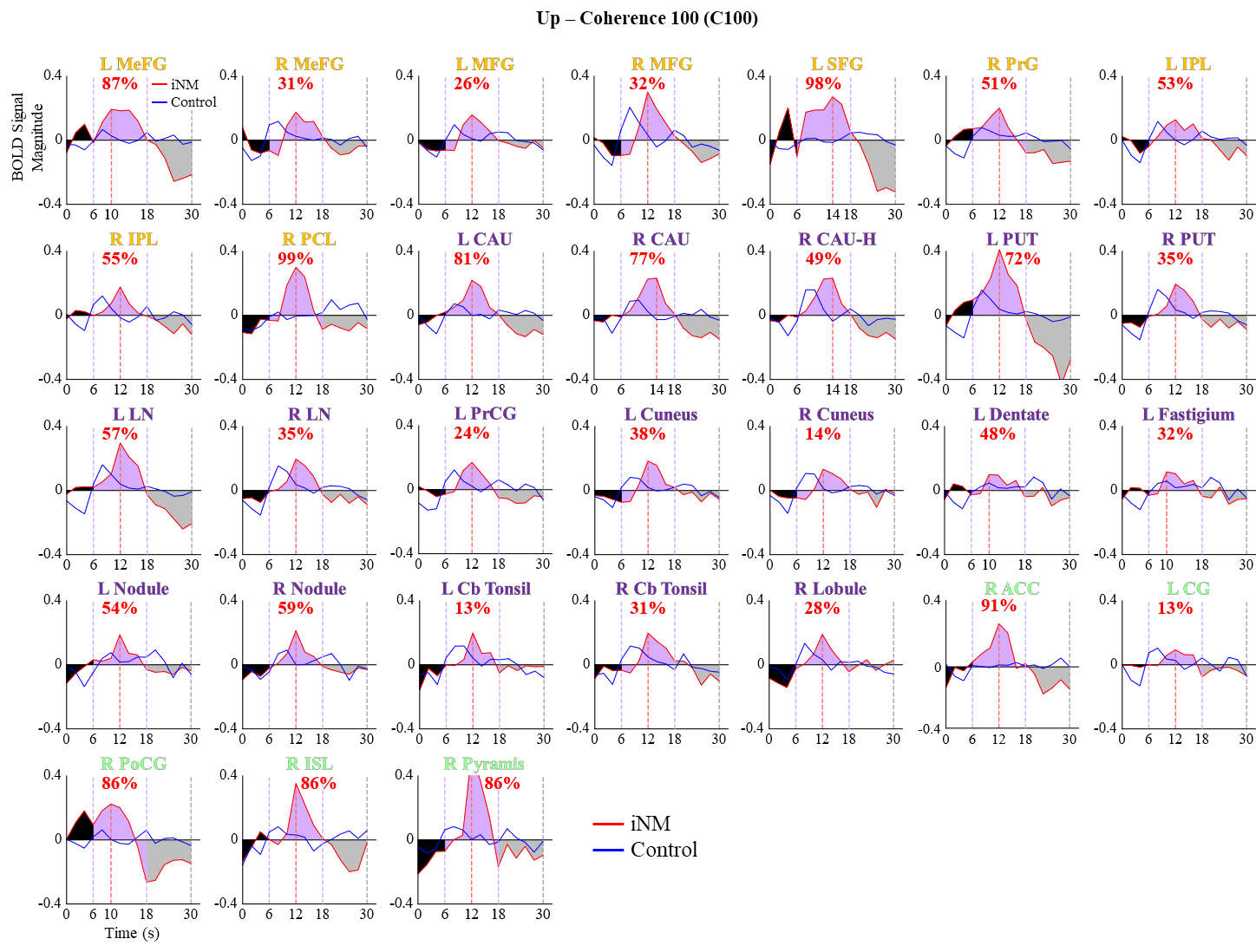
**Figure SI-5.** BOLD signal intensity of iNM (red) and control (blue) conditions are plotted as a function of time for all individual coherence level C100 regions in the up-motion direction. Each plot contains three sections: 1) a hemodynamic lag block (0-6 seconds); 2) a coherent motion block that was truncated to 18 seconds, as the signal intensity decreased across all ROIs (6-18 seconds) most likely due to participant fatigue in the task; and 3) a baseline-random motion block (18-30 seconds). The area under the BOLD signal intensity curve of the iNM condition was colored: black, hemodynamic lag; purple, coherent motion; and gray, baseline-random motion blocks. Percent change in area under the curve between control and iNM conditions for the coherent motion block is shown in red. Color of the individual region names indicate the network region, as shown under SI-1.

*The list of abbreviated regions include*: MeFG: medial frontal gyrus; MFG: middle frontal gyrus; SFG: superior frontal gyrus; FG: fusiform gyrus; INS: insula; IFG: inferior frontal gyrus; STG: superior temporal gyrus; PrCG: precentral gyrus; PC: posterior cingulate; PCL: paracentral lobule; CAU: caudate; CAU-H: caudate-head; PUT: putamen; LN: lentiform nucleus; Cb Tonsil: cerebellar tonsil; ACC: anterior cingulate cortex; ISL: Inferior Semilunar Lobule; CG: Cingulate Gyrus; PoCG: postcentral gyrus.

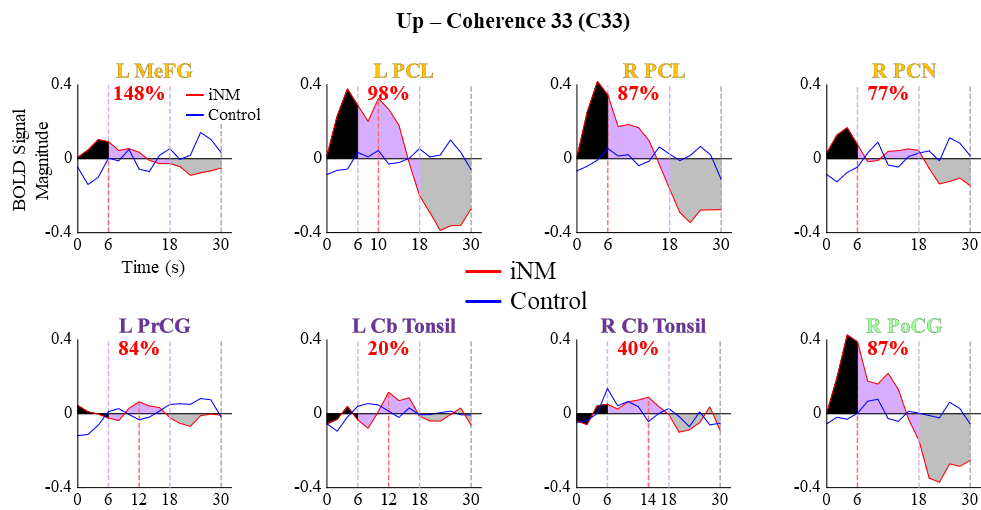

**Figure SI-6.** BOLD signal intensity of iNM (red) and control (blue) conditions are plotted as a function of time for all individual regions for coherence level C33 in the up-motion direction. Each plot contains three sections: 1) a hemodynamic lag block (0-6 seconds); 2) a coherent motion block that was truncated to 18 seconds, as the signal intensity decreased across all ROIs (6-18 seconds) most likely due to participant fatigue in the task; and 3) a baseline-random motion block (18-30 seconds). The area under the BOLD signal intensity curve of the iNM condition was colored: black, hemodynamic lag; purple, coherent motion; or gray, baseline-random motion blocks. The percent change of the area under the curve between control and iNM conditions for the coherent motion block is shown in red. Color of the individual region names indicate the region network.

*The list of abbreviated regions include*: MeFG: medial frontal gyrus; PCL: paracentral Lobule; PCN: precuneus; PrCG: precentral gyrus; Cb Tonsil: cerebellar tonsil; PoCG: postcentral gyrus;

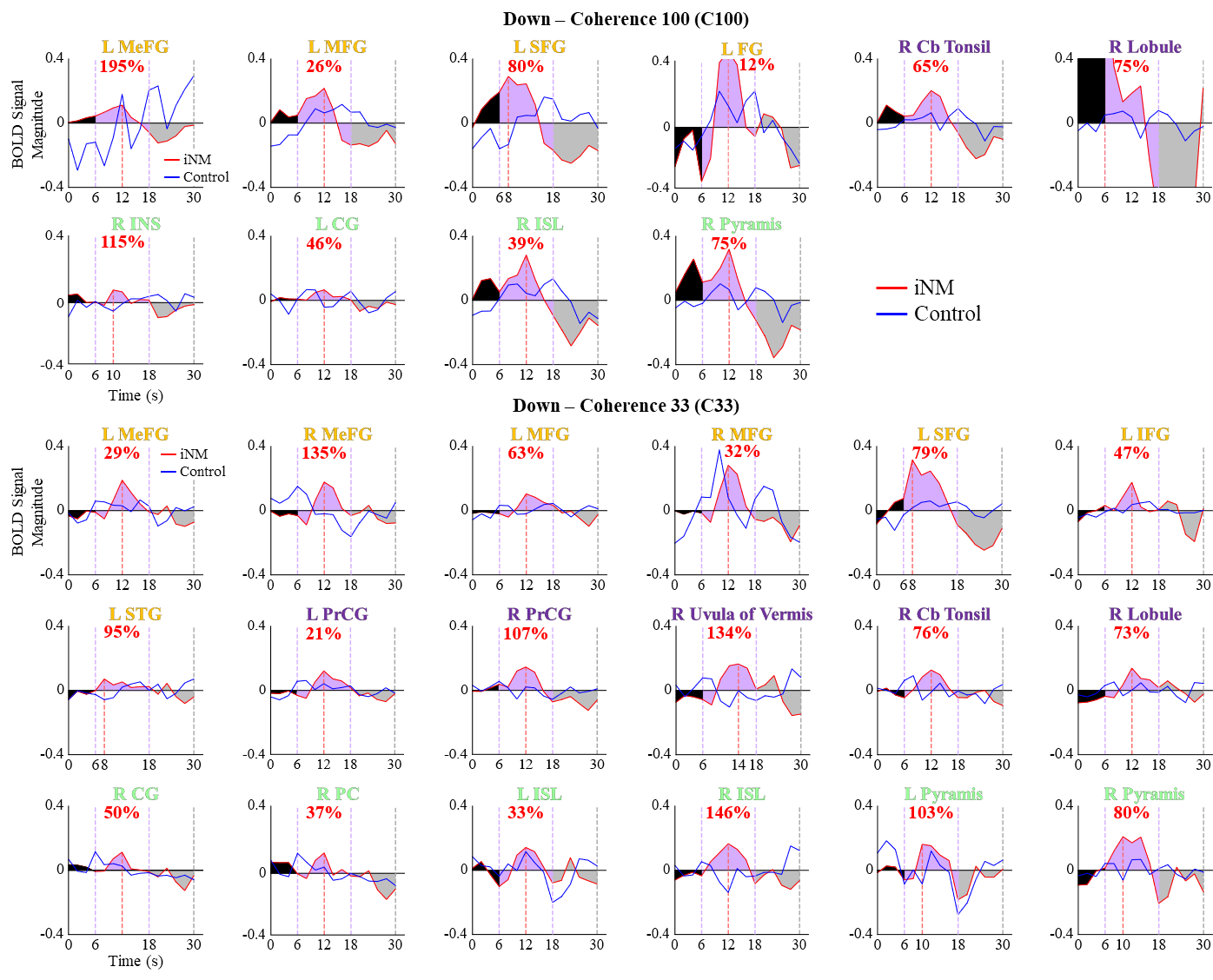
**Figure SI-7.** BOLD signal intensity of iNM (red) and control (blue) conditions are plotted as a function of time for all individual regions and coherence levels in the down-motion direction. Each plot contains three sections: 1) a hemodynamic lag block (0-6 seconds), 2) a coherent motion block that was truncated to 18 seconds, as the signal intensity plummeted across all ROIs due to potential participant fatigue in the task (6-18 seconds), and 3) a baseline-random motion block (18-30 seconds). The area under the BOLD signal intensity curve of the iNM condition is colored: black, hemodynamic lag; purple, coherent motion; or gray, baseline-random motion blocks. Percent change of the area under the curve between the control and iNM conditions for the coherent motion block is shown in red. Color of individual region names indicates the region network.

*The list of abbreviated regions include*: MeFG: medial frontal gyrus; MFG: middle frontal gyrus; SFG: superior frontal gyrus, IFG: inferior frontal gyrus; Cb Tonsil: cerebellar tonsil; R Lobule; INS: insula; CG: cingulate gyrus; ISL: inferior semilunar lobule; STG: superior temporal gyrus; PrCG: precentral gyrus; Cb tonsil: cerebellar tonsil; PC: posterior cingulate.

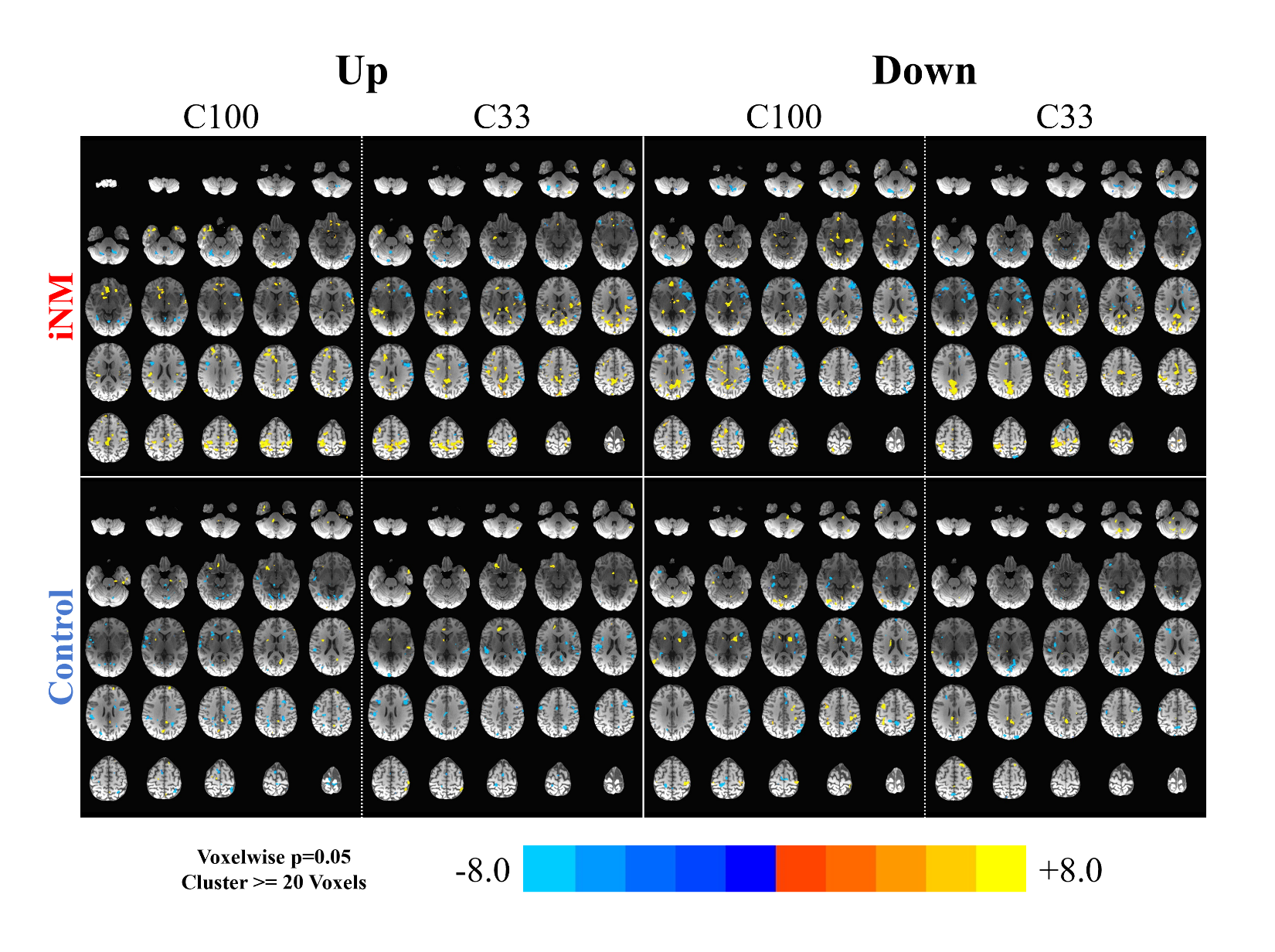
**Figure SI-8.** SVM-generated activation maps for each coherence level and motion direction in iNM (top) and control – no iNM (bottom) conditions: voxelwise p value = 0.05.

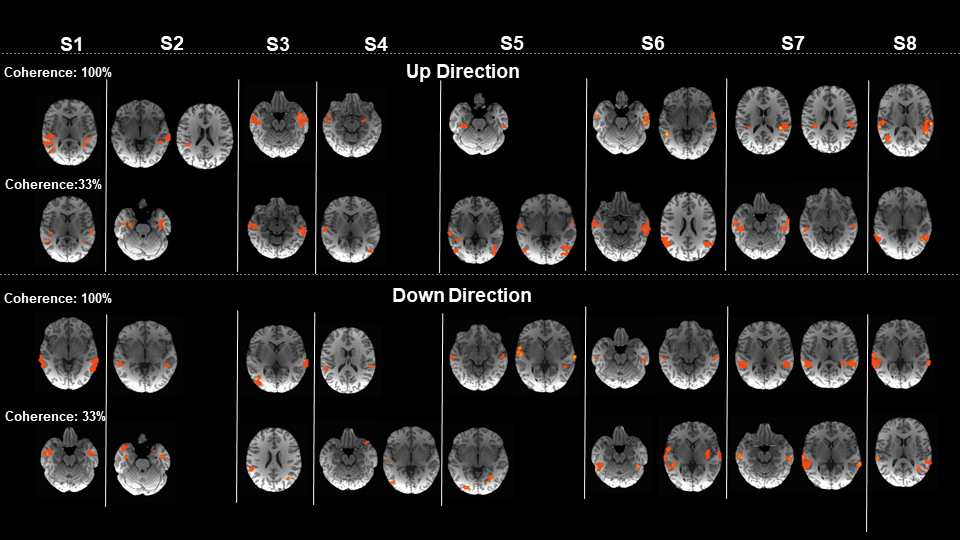

**Figure SI-9.** Individualized MT and MST anatomical and functional networks for each participant for Up direction at 100% coherence, 33% of coherence, and Down direction at 100% coherence and 33% coherence (p<0.05).

**Table S1a.** Peaks of activated regions generated by 3dExtrema for 100% Coherence Up Direction, control condition.

| TT_Daemon Atlas Region | Cluster | Cluster Size  (# of Voxels) | Region Size (# of Voxels) | RAI Peak Coordinates (mm) | | | Peak  Z-score |
| --- | --- | --- | --- | --- | --- | --- | --- |
|  |  |  |  | x | y | z |  |
| R central - lobules II-III | 1 | 67 | 682 | -11 | 47 | -19 | 7.738 |
| R superior & inferior semilunar |  | 7 | 153 | -35 | 65 | -25 | 6.397 |
| L declive - lobule VI |  | 38 | 636 | 5 | 59 | -16 | 5.958 |
| R inferior semilunar lobule VI |  | 67 | 682 | -26 | 59 | -25 | 5.422 |
| R tonsil |  | 84 | 526 | -47 | 50 | -37 | 5.331 |
| L uvula - lobule IX |  | 63 | 524 | 8 | 56 | -31 | 5.325 |
| L culmen - lobule IV-V |  | 38 | 636 | 2 | 50 | -4 | 4.861 |
| L putamen | 2 | 70 | 286 | 26 | -5 | 9 | 5.786 |
| L inferior frontal - pars triangularis | 3 | 64 | 1125 | 35 | -35 | 9 | 7.623 |
| L cingulate |  | 6 | 983 | 23 | -20 | 27 | 5.560 |
| L middle frontal gyrus |  | 67 | 1895 | 38 | -35 | -4 | 4.174 |
| L precentral (M1 - BA4) | 4 | 85 | 986 | 29 | 14 | 60 | 5.522 |
| R. putamen | 5 | 44 | 282 | -20 | -2 | -1 | 5.373 |
| R caudate |  | 12 | 160 | -8 | -5 | 3 | 3.707 |
| R anterior cingulate |  | 2 | 420 | -20 | -26 | 15 | 3.424 |
| L inferior parietal (BA40) | 6 | 79 | 790 | 41 | 38 | 45 | 5.336 |
| R precentral (M1 - BA4) | 7 | 83 | 1002 | -29 | 20 | 57 | 6.083 |
| L culmen - lobule IV-V | 8 | 38 | 636 | 35 | 32 | -31 | 5.323 |
| R precentral (M1 - BA4) | 9 | 83 | 1002 | -53 | -11 | 9 | 5.050 |
| L supplementary motor area | 10 | 8 | 999 | 2 | -5 | 48 | 3.932 |
| L superior frontal (BA6) | 11 | 8 | 999 | 17 | -11 | 48 | 3.756 |
| L superior frontal (BA8) |  | 21 | 1469 | 11 | -26 | 51 | 3.274 |
| L superior frontal (BA9) |  | 21 | 1469 | 14 | -17 | 54 | 3.002 |
| L pre-motor eye fields (BA8) | 12 | 85 | 986 | 47 | 2 | 36 | 3.736 |
| R pre-motor eye fields (BA8) | 13 | 26 | 1883 | -53 | -5 | 42 | 5.377 |
| L insula | 14 | 64 | 1125 | 38 | -23 | 9 | 3.688 |
| R superior frontal (BA6) | 15 | 6 | 1474 | -26 | 8 | 66 | 6.461 |
| R precentral (M1 - BA4/6) | 16 | 19 | 1155 | -50 | -2 | 24 | 3.262 |
| L inferior parietal (BA40) | 27 | 79 | 790 | 41 | 41 | 27 | 5.131 |
| R superior & inferior semilunar – lobules VIIa &VIIb | 18 | 14 | 218 | -35 | 65 | -40 | 6.656 |
| L superior & inferior semilunar | 19 | 63 | 524 | 41 | 50 | -37 | 5.256 |
| L inferior & middle temporal | 20 | 3 | 306 | 56 | 41 | -16 | 3.935 |
| L central - lobules II-III | 21 | 38 | 636 | 14 | 47 | -16 | 3.456 |
| R medial frontal | 22 | 24 | 1034 | -17 | -26 | 30 | 5.308 |
| R anterior cingulate |  | 2 | 420 | -17 | -23 | 18 | 3.751 |
| R superior & inferior semilunar – lobules VIIa &VIIb | 23 | 14 | 247 | -47 | 74 | -28 | 4.941 |
| L pre-motor eye fields (BA8) | 24 | 67 | 1895 | 29 | -2 | 42 | 6.596 |
| R postcentral (S1 - BA3/1/2) | 25 | 16 | 799 | -53 | 29 | 45 | 3.095 |

*Note: Regions may appear multiple times due to their inclusion in multiple clusters.*

**Table S1b.** Peaks of activated regions generated by 3dExtrema for 100% Coherence Up Direction, iNM condition.

| TT_Daemon Atlas Region | Cluster | Cluster Size  (# of Voxels) | Region Size (# of Voxels) | RAI Peak Coordinates (mm) | | | Peak  Z-score |
| --- | --- | --- | --- | --- | --- | --- | --- |
|  |  |  |  | x | y | z |  |
| R culmen and quadrangular - lobules IV-V | 1 | 20 | 682 | -14 | 38 | -25 | 5.669 |
| L superior & inferior semilunar |  | 27 | 524 | 14 | 56 | -37 | 5.645 |
| R biventral |  | 108 | 526 | -2 | 53 | -34 | 4.836 |
| R central lobule & central |  | 108 | 526 | -20 | 38 | -28 | 4.237 |
| R superior & inferior semilunar - lobule VI | 2 | 21 | 247 | -53 | 50 | -28 | 5.591 |
| R tuber & declive |  | 21 | 247 | -50 | 68 | -25 | 4.712 |
| R superior & inferior semilunar |  | 20 | 682 | -41 | 41 | -31 | 4.694 |
| R pyramis |  | 9 | 271 | -50 | 65 | -34 | 3.235 |
| L putamen | 3 | 5 | 57 | 11 | -8 | -10 | 6.989 |
| R nucleus accumbens |  | 6 | 420 | -2 | -2 | -7 | 5.457 |
| L caudate |  | 14 | 165 | 2 | -11 | 3 | 4.957 |
| R lingula - lobule I | 4 | 20 | 682 | -5 | 47 | -10 | 6.874 |
| L superior and inferior semilunar |  | 4 | 18 | 8 | 53 | -22 | 3.604 |
| R quadrangular |  | 20 | 682 | -14 | 47 | -10 | 2.741 |
| L middle occipital | 5 | 27 | 642 | 35 | 74 | -1 | 5.283 |
| L thalamus - medial dorsal nucleus | 6 | 0 | 11 | 5 | 17 | -13 | 5.854 |
| L red nucleus |  | 0 | 286 | 5 | 23 | -7 | 4.464 |
| L thalamus - proper | 7 | 27 | 642 | 47 | 65 | -4 | 4.643 |
| R middle temporal | 8 | 13 | 1042 | -20 | 50 | 51 | 5.418 |
| R inferior parietal |  | 3 | 799 | -41 | 38 | 48 | 3.726 |
| L superior & inferior semilunar | 9 | 42 | 636 | 38 | 32 | -28 | 4.728 |
| L culmen |  | 42 | 636 | 32 | 38 | -22 | 4.335 |
| L postcentral (BA3/1/2) | 10 | 2 | 725 | 32 | 26 | 42 | 8.063 |
| R medial frontal (BA10) | 11 | 15 | 1009 | -2 | -65 | -4 | 4.940 |
| L precentral (premotor BA6) | 12 | 11 | 1895 | 23 | 14 | 60 | 3.335 |
| R medial frontal (BA10) | 13 | 16 | 983 | 20 | -2 | 48 | 4.063 |
| L hippocampus | 14 | 14 | 165 | 35 | 32 | -7 | 5.313 |
| L posterior cingulate | 15 | 0 | 239 | 5 | 38 | 15 | 3.686 |
| L hippocampus |  | 5 | 465 | 17 | 38 | 6 | 3.163 |
| R precentral (premotor BA6) | 16 | 25 | 1002 | -32 | 14 | 54 | 4.100 |
| R supplementary motor (BA6) | 17 | 15 | 1009 | -5 | 2 | 54 | 3.549 |
| R superior temporal (BA38) | 18 | 8 | 1288 | -41 | -8 | -22 | 5.008 |
| R posterior cingulate | 19 | 3 | 230 | -11 | 41 | 15 | 3.967 |
| L parahippocampal | 20 | 5 | 465 | 17 | 17 | -22 | 4.749 |
| L middle cingulate | 21 | 16 | 983 | 11 | 26 | 33 | 3.685 |
| R posterior parahippocampal | 22 | 2 | 202 | -2 | 38 | 66 | 7.883 |
| R superior parietal | 23 | 13 | 751 | -29 | 38 | 60 | 4.127 |
| R precentral (premotor BA6) | 24 | 25 | 1002 | -29 | 23 | 63 | 6.241 |
| L superior parietal | 25 | 12 | 999 | 14 | 50 | 45 | 7.719 |
| R putamen | 26 | 0 | 45 | -26 | -17 | 3 | 3.593 |
| R middle cingulate | 27 | 10 | 1034 | -14 | 8 | 36 | 3.857 |
| R precentral (premotor BA6) | 28 | 10 | 1883 | -26 | 14 | 60 | 4.427 |
| L precentral (premotor BA6) | 29 | 12 | 999 | 5 | 26 | 69 | 4.296 |

*Note: Regions may appear multiple times due to their inclusion in multiple clusters.*

**Table S2a.** Peaks of activated regions generated by 3dExtrema for 100% Coherence Down Direction, control condition.

| TT_Daemon Atlas Region | Cluster | Cluster Size  (# of Voxels) | Region Size (# of Voxels) | RAI Peak Coordinates (mm) | | | Peak  Z-score |
| --- | --- | --- | --- | --- | --- | --- | --- |
|  |  |  |  | x | y | z |  |
| R inferior semilunar - lobule VIIb | 1 | 82 | 526 | -35 | 65 | -46 | 8.066 |
| R culmen - lobules IV-V |  | 49 | 218 | -17 | 62 | -40 | 6.213 |
| R quadrangular - lobule III |  | 82 | 526 | -32 | 59 | -37 | 4.377 |
| L superior parietal | 2 | 45 | 999 | 20 | 65 | 51 | 4.332 |
| L precentral (M1) | 3 | 52 | 1895 | 29 | 8 | 63 | 6.004 |
| L inferior frontal (BA9) | 4 | 22 | 1125 | 56 | -8 | 33 | 4.354 |
| L premotor area, frontal eye fields (BA8) |  | 35 | 986 | 38 | 2 | 42 | 2.594 |
| L culmen - lobules IV-V | 5 | 0 | 636 | 50 | 47 | -22 | 3.712 |
| L inferior temporal (BA37) |  | 18 | 306 | 53 | 47 | -13 | 3.584 |
| L inferior semilunar - lobule VIIb | 6 | 10 | 219 | 23 | 65 | -40 | 8.497 |
| R precentral (M1 - BA6) | 7 | 13 | 1155 | -59 | -8 | 24 | 3.586 |
| L rolandic operculum | 8 | 11 | 790 | 38 | 38 | 27 | 4.121 |
| L postcentral (S1 - BA3/1/2) |  | 13 | 725 | 35 | 29 | 39 | 3.119 |
| L middle temporal (BA39) | 9 | 46 | 1233 | 47 | 56 | 12 | 4.082 |
| L inferior temporal (BA37) | 10 | 13 | 367 | 53 | 11 | -25 | 4.590 |
| L inferior frontal - pars triangularis | 11 | 22 | 1125 | 47 | -29 | 15 | 3.002 |
| L middle frontal (BA46) |  | 52 | 1895 | 47 | -38 | 15 | 2.820 |
| L superior medial (BA9) | 12 | 27 | 1469 | 14 | -65 | 21 | 4.779 |
| L culmen - lobules IV-V | 13 | 34 | 524 | 20 | 47 | -31 | 5.615 |
| L declive - lobule VI |  | 0 | 28 | 17 | 56 | -28 | 3.200 |
| L middle temporal (BA21) | 14 | 46 | 1233 | 65 | 44 | -4 | 4.251 |

*Note: Regions may appear multiple times due to their inclusion in multiple clusters.*

**Table S2b.** Peaks of activated regions generated by 3dExtrema for 100% Coherence Down Direction, iNM condition.

| TT_Daemon Atlas Region | Cluster | Cluster Size  (# of Voxels) | Region Size (# of Voxels) | RAI Peak Coordinates (mm) | | | Peak  Z-score |
| --- | --- | --- | --- | --- | --- | --- | --- |
|  |  |  |  | x | y | z |  |
| R inferior semilunar - lobule VIIb | 1 | 119 | 526 | -44 | 53 | -37 | 10.208 |
| R superior semilunar - lobule VIIa |  | 9 | 682 | -50 | 44 | -28 | 5.676 |
| R pyramis - lobule VIII |  | 30 | 271 | -20 | 83 | -34 | 4.786 |
| L inferior frontal - pars orbital | 2 | 19 | 1125 | 29 | -26 | -4 | 4.153 |
| L inferior frontal - pars triangularis |  | 19 | 1125 | 38 | -35 | 3 | 3.442 |
| L caudate |  | 0 | 165 | 20 | -20 | 3 | 3.124 |
| L lingual (V1 - BA18) | 3 | 10 | 555 | 17 | 98 | -4 | 4.224 |
| L cuneus (V1 - BA18) |  | 9 | 704 | 26 | 95 | -1 | 3.429 |
| L precuneus (V1 - BA17) |  | 9 | 704 | 20 | 86 | 9 | 3.085 |
| L anterior middle cingulate | 4 | 14 | 983 | 14 | -20 | 27 | 4.175 |
| L anterior cingulate (BA32) |  | 15 | 393 | 11 | -14 | 21 | 4.049 |
| L medial frontal (BA9) |  | 8 | 1895 | 26 | -23 | 27 | 3.988 |
| L inferior frontal - pars triangularis |  | 8 | 1895 | 32 | -17 | 21 | 3.146 |
| L precentral (M1 - BA6) | 5 | 10 | 1002 | -14 | 20 | 69 | 4.848 |
| R middle temporal | 6 | 0 | 160 | -38 | 17 | -10 | 6.916 |
| R parahippocampal |  | 8 | 484 | -38 | 29 | -10 | 2.734 |
| R middle occipital (V2) | 7 | 0 | 649 | -35 | 56 | 3 | 6.586 |
| R inferior temporal (BA20) |  | 5 | 370 | -32 | 53 | -10 | 3.818 |
| R declive - lobule VI |  | 9 | 682 | -26 | 50 | -16 | 2.911 |
| R precentral (M1 - BA3) | 8 | 7 | 1009 | -5 | 26 | 66 | 3.459 |
| L parahippocampal | 9 | 7 | 465 | 38 | 35 | -7 | 4.572 |
| L middle temporal |  | 0 | 1233 | 44 | 29 | -7 | 4.334 |
| L supplementary eye fields (BA6) | 10 | 17 | 999 | 14 | 2 | 51 | 4.995 |
| L superior medial frontal | 11 | 15 | 393 | 11 | -50 | -1 | 3.812 |
| L anterior cingulate |  | 15 | 393 | 2 | -44 | 3 | 3.157 |
| L thalamus - pulvinar | 12 | 1 | 286 | 20 | 35 | 3 | 4.534 |
| L superior parietal | 13 | 11 | 230 | 23 | 68 | 54 | 3.025 |
| L superior frontal (BA10) | 14 | 16 | 1469 | 23 | -59 | 18 | 4.180 |
| L superior medial frontal | 15 | 16 | 1469 | 2 | -32 | 51 | 3.461 |
| L paracentral | 16 | 8 | 986 | 17 | 23 | 63 | 3.763 |
| R parahippocampal | 17 | 0 | 160 | -35 | 32 | -7 | 2.751 |
| L thalamus - pulvinar | 18 | 7 | 286 | 17 | 11 | -7 | 4.112 |
| L lentiform nucleus |  | 7 | 286 | 17 | 5 | -4 | 3.075 |
| L parahippocampal | 19 | 7 | 465 | 14 | 20 | -7 | 4.668 |

*Note: Regions may appear multiple times due to their inclusion in multiple clusters.*

**Table S3a.** Peaks of activated regions generated by 3dExtrema for 33% Coherence Up Direction, control condition.

| TT_Daemon Atlas Region | Cluster | Cluster Size  (# of Voxels) | Region Size (# of Voxels) | RAI Peak Coordinates (mm) | | | Peak  Z-score |
| --- | --- | --- | --- | --- | --- | --- | --- |
|  |  |  |  | x | y | z |  |
| R claustrum | 1 | 1 | 45 | -26 | -23 | 3 | 7.808 |
| R inferior frontal - operculum |  | 131 | 1155 | -59 | -17 | -1 | 6.307 |
| R precentral (primary motor - BA4) |  | 131 | 1155 | -59 | -20 | 6 | 4.740 |
| R insula |  | 131 | 1155 | -32 | -26 | -4 | 4.262 |
| R inferior frontal - orbital |  | 131 | 1155 | -32 | -26 | -10 | 3.314 |
| R postcentral (primary sensory) | 2 | 16 | 751 | -11 | 38 | 66 | 5.995 |
| L paracentral (BA3/1/2) |  | 4 | 999 | 8 | 23 | 66 | 5.259 |
| R superior parietal |  | 5 | 1042 | -11 | 50 | 63 | 3.965 |
| L precentral (primary motor) |  | 34 | 986 | 11 | 17 | 66 | 3.817 |
| R middle frontal (BA10) | 3 | 64 | 1883 | -44 | -50 | 6 | 10.178 |
| R superior frontal - orbital |  | 34 | 1474 | -32 | -59 | -7 | 3.072 |
| L inferior frontal – pars triangularis | 4 | 48 | 1895 | 50 | -38 | -4 | 5.323 |
| L middle frontal |  | 48 | 1895 | 44 | -50 | 12 | 3.899 |
| R superior frontal | 5 | 64 | 1883 | -23 | 2 | 57 | 4.291 |
| R precentral (primary motor - BA4) |  | 31 | 1002 | -32 | 17 | 54 | 4.067 |
| L claustrum | 6 | 8 | 52 | 29 | -14 | -1 | 5.680 |
| L middle frontal |  | 0 | 393 | 23 | -38 | 3 | 5.428 |
| L insula |  | 11 | 515 | 32 | -20 | -1 | 4.160 |
| L caudate |  | 0 | 165 | 20 | -26 | 3 | 3.453 |
| L culmen - lobules IV-V | 7 | 41 | 524 | 32 | 53 | -34 | 8.354 |
| R posterior cingulate | 8 | 0 | 1034 | -26 | 41 | 24 | 5.114 |
| R precuneus |  | 0 | 1034 | -26 | 50 | 24 | 3.245 |
| R inferior frontal - operculum | 9 | 131 | 1155 | -62 | -14 | 21 | 10.199 |
| R middle temporal | 10 | 13 | 1244 | -47 | 35 | -4 | 5.257 |
| L precentral (primary motor) | 11 | 34 | 986 | 29 | 17 | 54 | 3.510 |
| R culmen - lobules IV-V | 12 | 28 | 526 | -26 | 53 | -31 | 6.746 |
| L inferior temporal | 13 | 6 | 367 | 53 | 35 | -19 | 4.614 |
| L middle temporal |  | 14 | 1233 | 62 | 32 | -13 | 4.370 |
| R supplementary motor area | 14 | 34 | 1474 | -11 | -8 | 60 | 4.373 |
| R inferior frontal – pars triangularis | 15 | 64 | 1883 | -56 | -32 | 15 | 7.716 |
| R middle temporal | 16 | 13 | 1244 | -44 | -8 | -28 | 4.186 |
| L lingual (primary visual - BA18) | 17 | 9 | 555 | 5 | 95 | -13 | 7.289 |
| L postcentral (primary motor - BA2) | 18 | 18 | 725 | 35 | 29 | 45 | 5.212 |
| L putamen | 19 | 9 | 286 | 29 | -5 | 3 | 5.176 |
| L middle Temporal | 20 | 14 | 1233 | 44 | 32 | -7 | 3.554 |
| L inferior frontal | 21 | 10 | 531 | -32 | -26 | 18 | 6.779 |
| L postcentral (primary motor - BA2) | 22 | 31 | 1002 | -29 | 29 | 45 | 3.499 |
| R posterior cingulate | 23 | 10 | 983 | 20 | 44 | 24 | 5.824 |
| R supplementary motor area | 24 | 34 | 1009 | -11 | 11 | 60 | 3.456 |
| R pyramis - lobule VIII | 25 | 0 | 218 | -44 | 65 | -43 | 3.511 |
| R cerebellar tonsil |  | 28 | 526 | -41 | 59 | -43 | 2.936 |
| R superior medial frontal (BA10) | 26 | 34 | 1009 | -14 | -65 | -4 | 4.104 |

*Note: Regions may appear multiple times due to their inclusion in multiple clusters.*

**Table S3b.** Peaks of activated regions generated by 3dExtrema for 33% Coherence Up Direction, iNM condition.

| TT_Daemon Atlas Region | Cluster | Cluster Size  (# of Voxels) | Region Size (# of Voxels) | RAI Peak Coordinates (mm) | | | Peak  Z-score |
| --- | --- | --- | --- | --- | --- | --- | --- |
|  |  |  |  | x | y | z |  |
| L superior parietal | 1 | 18 | 230 | 20 | 65 | 60 | 5.051 |
| R precuneus (BA7) |  | 69 | 1042 | -11 | 62 | 51 | 4.634 |
| R precentral (primary motor - BA6) | 2 | 25 | 1002 | -41 | 5 | 51 | 5.673 |
| R superior frontal |  | 58 | 1883 | -23 | 2 | 60 | 4.214 |
| R supplementary motor area |  | 12 | 1034 | -14 | -2 | 48 | 3.245 |
| L middle temporal | 3 | 15 | 1233 | 32 | 65 | 18 | 5.140 |
| L superior frontal | 4 | 10 | 983 | 20 | -8 | 45 | 5.247 |
| R ventral dentate nucleus | 5 | 24 | 526 | -11 | 44 | -34 | 5.862 |
| R superior & inferior semilunar |  | 2 | 682 | -17 | 44 | -25 | 3.976 |
| L superior medial frontal | 6 | 35 | 1469 | 20 | -62 | 24 | 4.111 |
| R globus pallidus | 7 | 24 | 282 | -14 | 8 | 3 | 3.942 |
| R putamen |  | 24 | 282 | -20 | -8 | -7 | 2.717 |
| R caudate |  | 3 | 160 | -8 | -5 | 3 | 2.578 |
| L paracentral | 8 | 21 | 999 | 5 | 14 | 69 | 6.374 |
| L precentral (primary motor) |  | 12 | 986 | 14 | 20 | 66 | 4.272 |
| L ventral cerebellar dentate | 9 | 29 | 524 | 20 | 56 | -34 | 3.000 |
| R paracentral | 10 | 25 | 1009 | -2 | 26 | 66 | 6.024 |
| R superior parietal | 11 | 69 | 1042 | -20 | 44 | 45 | 5.461 |
| R parahippocampal | 12 | 5 | 484 | -23 | 20 | -25 | 5.682 |
| R lingual | 13 | 2 | 682 | -2 | 38 | -10 | 4.367 |
| L middle frontal (BA10) | 14 | 43 | 1895 | 35 | -53 | 9 | 3.514 |
| L inferior frontal - pars triangularis |  | 8 | 1125 | 44 | -38 | 9 | 3.471 |
| L superior medial frontal | 15 | 21 | 999 | 8 | -29 | 42 | 6.706 |
| R superior & inferior semilunar | 16 | 29 | 524 | 44 | 47 | -37 | 4.318 |
| R thalamus - prefrontal | 17 | 1 | 284 | -8 | 17 | -7 | 5.130 |
| R entorhinal cortex | 18 | 7 | 157 | -32 | 2 | -28 | 7.551 |
| L posterior cingulate | 19 | 10 | 983 | 20 | 41 | 27 | 3.508 |
| R superior frontal | 20 | 25 | 1009 | -11 | 14 | 63 | 5.777 |
| R inferior parietal | 21 | 8 | 799 | -41 | 41 | 48 | 3.595 |
| R supplementary motor area | 22 | 25 | 1009 | -8 | 5 | 60 | 3.463 |
| R inferior frontal – pars triangularis | 23 | 10 | 1155 | -56 | -29 | -1 | 5.183 |

*Note: Regions may appear multiple times due to their inclusion in multiple clusters.*

**Table S4a.** Peaks of activated regions generated by 3dExtrema for 33% Coherence Down Direction, control condition.

| TT_Daemon Atlas Region | Cluster | Cluster Size  (# of Voxels) | Region Size (# of Voxels) | RAI Peak Coordinates (mm) | | | Peak  Z-score |
| --- | --- | --- | --- | --- | --- | --- | --- |
|  |  |  |  | x | y | z |  |
| L precentral (primary motor) | 1 | 69 | 986 | 32 | 8 | 39 | 9.833 |
| L inferior frontal - operculum |  | 60 | 1125 | 47 | -8 | 27 | 3.976 |
| L biventral | 2 | 36 | 219 | 41 | 68 | -37 | 10.117 |
| L superior frontal - orbital | 3 | 47 | 1469 | 23 | -53 | -4 | 4.120 |
| L superior medial |  | 5 | 393 | 17 | -50 | -1 | 4.015 |
| L inferior temporal | 4 | 19 | 367 | 47 | 38 | -19 | 5.489 |
| L superior frontal - orbital | 5 | 47 | 1469 | 38 | -56 | 15 | 4.806 |
| L inferior frontal – pars triangularis |  | 111 | 1895 | 41 | -53 | 9 | 4.514 |
| L middle temporal | 6 | 47 | 306 | 56 | 53 | -4 | 6.391 |
| L supramarginal | 7 | 19 | 199 | 41 | 44 | 33 | 4.388 |
| L inferior frontal – pars triangularis | 8 | 60 | 1125 | 50 | -32 | 12 | 3.938 |
| R flocculus - lobule IX | 9 | 44 | 526 | -11 | 59 | -43 | 5.471 |
| R biventral - lobule VIII |  | 44 | 526 | -23 | 56 | -40 | 4.391 |
| R posterior cingulate | 10 | 18 | 1034 | -11 | 29 | 24 | 6.579 |
| L inferior semilunar | 11 | 19 | 243 | 38 | 62 | -28 | 4.161 |
| L precentral (primary motor) | 12 | 111 | 1895 | 38 | -20 | 33 | 6.316 |
| R supramarginal | 13 | 10 | 202 | -35 | 53 | 36 | 3.770 |
| R superior semilunar | 14 | 20 | 682 | -47 | 44 | -28 | 4.910 |
| R precentral | 15 | 10 | 1002 | -17 | 20 | 66 | 6.150 |
| L inferior semilunar - lobule VIII | 16 | 61 | 524 | 17 | 62 | -31 | 5.687 |
| R Inferior temporal | 17 | 19 | 315 | -59 | 26 | -22 | 4.859 |
| R superior semilunar - lobule VII | 18 | 3 | 247 | -32 | 59 | -28 | 5.981 |
| L inferior parietal | 19 | 6 | 230 | 38 | 62 | 51 | 4.200 |
| L precentral (primary motor) | 20 | 69 | 986 | 29 | 20 | 54 | 4.534 |
| R lingual - lobule I | 21 | 44 | 526 | -14 | 44 | -37 | 3.752 |
| L nodulus - lobule X | 22 | 61 | 524 | 17 | 35 | -37 | 6.591 |
| R inferior semilunar - lobule VII | 23 | 10 | 271 | -17 | 71 | -28 | 6.228 |
| L middle temporal | 24 | 38 | 1233 | 47 | 32 | -7 | 4.113 |
| R inferior frontal - orbital | 25 | 24 | 1883 | -44 | -38 | -13 | 5.294 |
| R superior frontal (BA10) | 26 | 17 | 1474 | -23 | -59 | 6 | 6.655 |
| L superior semilunar - lobule VII | 27 | 61 | 524 | 35 | 41 | -40 | 5.429 |
| R postcentral (primary sensory) | 28 | 14 | 751 | -14 | 44 | 66 | 6.824 |
| R middle temporal | 29 | 19 | 315 | -62 | 53 | -7 | 5.051 |
| R inferior frontal – pars triangularis | 30 | 12 | 1155 | -56 | -38 | 3 | 3.876 |
| R inferior frontal – pars triangularis | 31 | 12 | 1155 | -23 | -29 | 3 | 3.278 |
| R medial frontal |  | 0 | 420 | -23 | -35 | 6 | 3.157 |
| R postcentral gyrus (primary sensory) | 32 | 12 | 799 | -38 | 32 | 30 | 3.630 |
| R biventral - lobule VIII | 33 | 44 | 526 | -44 | 59 | -43 | 4.067 |
| R precuneus (BA7) | 34 | 3 | 1042 | -5 | 77 | 48 | 3.887 |
| R inferior frontal - operculum | 35 | 24 | 1883 | -50 | -14 | 36 | 2.921 |
| R precentral (primary motor) |  | 10 | 1002 | -41 | -8 | 36 | 2.733 |

*Note: Regions may appear multiple times due to their inclusion in multiple clusters.*

**Table S4b.** Peaks of activated regions generated by 3dExtrema for 33% Coherence Down Direction, iNM condition.

| TT_Daemon Atlas Region | Cluster | Cluster Size  (# of Voxels) | Region Size (# of Voxels) | RAI Peak Coordinates (mm) | | | Peak  Z-score |
| --- | --- | --- | --- | --- | --- | --- | --- |
|  |  |  |  | x | y | z |  |
| R inferior semilunar - lobule VII | 1 | 118 | 218 | -38 | 74 | -40 | 11.290 |
| R tuber - lobule VII |  | 50 | 271 | -32 | 80 | -31 | 9.368 |
| R biventral - lobule VIII |  | 118 | 218 | -5 | 74 | -40 | 7.628 |
| R tonsil |  | 27 | 247 | -44 | 77 | -28 | 5.187 |
| R superior semilunar - lobule VII |  | 167 | 526 | -32 | 50 | -31 | 4.716 |
| R culmen - lobules IV-V |  | 12 | 682 | -53 | 41 | -28 | 4.536 |
| R uvula |  | 50 | 271 | -5 | 86 | -28 | 3.672 |
| L posterior cingulate (BA31) | 2 | 69 | 983 | 23 | 20 | 33 | 6.678 |
| L precentral (primary motor) |  | 11 | 986 | 20 | 17 | 57 | 5.433 |
| L middle cingulate (BA24) |  | 69 | 983 | 23 | 17 | 45 | 5.118 |
| L insula |  | 2 | 515 | 26 | 14 | 21 | 4.848 |
| R middle cingulate (BA24) | 3 | 23 | 1034 | -23 | 32 | 42 | 7.278 |
| R postcentral (primary sensory) |  | N/A | N/A | -29 | 23 | 36 | 6.169 |
| R inferior parietal |  | 0 | 799 | -29 | 23 | 30 | 5.127 |
| R postcentral (primary sensory) |  | 12 | 160 | -23 | 29 | 21 | 3.452 |
| L posterior cingulate (BA31) | 4 | 6 | 239 | 2 | 38 | 18 | 4.583 |
| L inferior semilunar | 5 | 7 | 219 | 29 | 65 | -43 | 6.051 |
| L biventral - lobule VIII |  | 24 | 524 | 29 | 59 | -46 | 5.504 |
| L paracentral (supplemental motor) | 6 | 38 | 999 | 5 | 5 | 57 | 4.366 |
| R precentral (primary motor) | 7 | 12 | 1883 | -26 | 14 | 60 | 3.813 |
| R paracentral (supplemental motor) |  | 15 | 1009 | -11 | 11 | 63 | 2.939 |
| L middle temporal | 8 | 20 | 306 | 53 | 26 | -22 | 6.363 |
| L inferior frontal - orbital | 9 | 24 | 1125 | 53 | -26 | -7 | 6.108 |
| L superior temporal |  | 16 | 1321 | 47 | -11 | -13 | 4.365 |
| L anterior cingulate | 10 | 12 | 160 | -2 | -5 | -1 | 3.764 |
| R caudate |  | 12 | 160 | -11 | -11 | 6 | 2.947 |
| L supramarginal | 11 | 1 | 199 | 35 | 38 | 33 | 4.734 |
| L superior parietal | 12 | 15 | 230 | 23 | 59 | 60 | 4.231 |
| L precuneus |  | 15 | 230 | 11 | 62 | 54 | 3.229 |
| L thalamus - proper | 13 | 3 | 286 | 14 | 32 | 9 | 3.342 |
| L hippocampus |  | 3 | 286 | 23 | 35 | 3 | 3.289 |
| R middle cingulate (BA24) | 14 | 23 | 1034 | -17 | 14 | 39 | 4.126 |
| R paracentral | 15 | 12 | 1002 | -11 | 20 | 69 | 6.555 |
| R supramarginal |  | 0 | 1474 | -11 | 14 | 72 | 3.909 |
| R supramarginal | 16 | 38 | 999 | 5 | 26 | 66 | 4.254 |
| L parahippocampal | 17 | 9 | 165 | 26 | -2 | -28 | 5.212 |
| L inferior frontal - operculum | 18 | 29 | 1895 | 38 | -23 | 30 | 4.897 |
| L precentral (primary motor) | 19 | 11 | 986 | 41 | -2 | 36 | 3.581 |
| L middle frontal (BA10) | 20 | 29 | 1895 | 23 | -62 | 9 | 3.291 |
| L primary motor (BA4) | 21 | 11 | 986 | 14 | 26 | 69 | 5.870 |
| R subcallosal (BA34) | 22 | 15 | 1009 | -8 | -5 | -16 | 5.306 |
| R middle temporal | 23 | 0 | 1244 | -47 | -5 | -16 | 4.302 |
| L supramarginal | 24 | 8 | 725 | 50 | 26 | 33 | 9.381 |
| L parahippocampal | 25 | 6 | 465 | 14 | 11 | -22 | 7.300 |
| L middle frontal | 26 | 69 | 983 | 23 | -17 | 33 | 3.931 |

*Note: Regions may appear multiple times due to their inclusion in multiple clusters.*

**Table S5a.** Support vector machine (SVM)-generated areas under the receiver-operator curve (AUC-ROC) performance values across all subjects as a function of different coherence levels and permutations of training-testing data types (control vs individualized neuromodulation [iNM]).

| Coherence – Direction | iNM AUROC | Control AUROC | p-value |
| --- | --- | --- | --- |
| C100 – Up | 0.68 | 0.52 | 0.001 |
| C33 – Up | 0.64 | 0.54 | 0.045 |
| C100 – Down | 0.71 | 0.53 | 0.004 |
| C33 - Down | 0.63 | 0.55 | 0.048 |

**Table S5b.** SVM-generated AUC-ROC performance values across all subjects as a function of up and down directions, coherence levels, and network involvement in the control and iNM conditions.

| Coherence - Direction | Network | iNM AUROC | Control AUROC | p-value |
| --- | --- | --- | --- | --- |
| C100 - Up | Visual Perception | 0.58 | 0.55 | 0.06 |
|  | Visual Imagery | 0.59 | 0.56 | 0.08 |
|  | Motor Planning | 0.63 | 0.55 | 0.05 |
|  | Selective Extero-Interoceptive Attention | 0.61 | 0.53 | 0.004 |
| C33 - Up | Visual Perception | 0.59 | 0.51 | 0.07 |
|  | Visual Imagery | 0.63 | 0.52 | 0.004 |
|  | Motor Planning | 0.63 | 0.50 | 0.02 |
|  | Selective Extero-Interoceptive Attention | 0.65 | 0.52 | 0.004 |
| C100 - Down | Visual Perception | 0.61 | 0.50 | 0.0006 |
|  | Visual Imagery | 0.64 | 0.54 | 0.02 |
|  | Motor Planning | 0.68 | 0.49 | 0.0007 |
|  | Selective Extero-Interoceptive Attention | 0.72 | 0.49 | 0.0008 |
| C33 - Down | Visual Perception | 0.56 | 0.56 | 0.32 |
|  | Visual Imagery | 0.62 | 0.54 | 0.29 |
|  | Motor Planning | 0.57 | 0.56 | 0.32 |
|  | Selective Extero-Interoceptive Attention | 0.61 | 0.60 | 0.86 |
